# Using Mechanistic Models and Machine Learning to Design Single-Color Multiplexed Nascent Chain Tracking Experiments

**DOI:** 10.1101/2023.01.25.525583

**Authors:** William S. Raymond, Sadaf Ghaffari, Luis U. Aguilera, Eric Ron, Tatsuya Morisaki, Zachary R. Fox, Michael P. May, Timothy J. Stasevich, Brian Munsky

## Abstract

mRNA translation is the ubiquitous cellular process of reading messenger-RNA strands into functional proteins. Over the past decade, large strides in microscopy techniques have allowed observation of mRNA translation at a single-molecule resolution for self-consistent time-series measurements in live cells. Dubbed Nascent chain tracking (NCT), these methods have explored many temporal dynamics in mRNA translation uncaptured by other experimental methods such as ribosomal profiling, smFISH, pSILAC, BONCAT, or FUNCAT-PLA. However, NCT is currently restricted to the observation of one or two mRNA species at a time due to limits in the number of resolvable fluorescent tags. In this work, we propose a hybrid computational pipeline, where detailed mechanistic simulations produce realistic NCT videos, and machine learning is used to assess potential experimental designs for their ability to resolve multiple mRNA species using a single fluorescent color for all species. Through simulation, we show that with careful application, this hybrid design strategy could in principle be used to extend the number of mRNA species that could be watched simultaneously within the same cell. We present a simulated example NCT experiment with seven different mRNA species within the same simulated cell and use our ML labeling to identify these spots with 90% accuracy using only two distinct fluorescent tags. The proposed extension to the NCT color palette should allow experimentalists to access a plethora of new experimental design possibilities, especially for cell signalling applications requiring simultaneous study of multiple mRNAs.

## 1 Introduction

mRNA translation is the process of reading messenger RNA strands to create functional proteins and is a crucial underpinning of known cellular life. With such a vital role, mRNA translation has been the subject of intense study over the past decades (Neelagandan et al., 2020; Das et al., 2021; Knight et al., 2020; Basyuk et al., 2020). Despite the focus, the effects that cellular signals have on the translation of individual mRNA molecules remains elusive due to two key factors: the staggering amount of dynamics, mechanisms, and modifications affecting the in vivo mRNA transcriptome heterogeneity and the limitations of experimental techniques that can accurately and informatively probe these dynamics molecule-by-molecule and in a time-resolved manner. Previous methods used to probe mRNA translation dynamics such as ribosomal footprinting (Brar and Weissman, 2015; Ingolia, 2016), RNA-seq (Wang et al., 2009; Stark et al., 2019; Chen et al., 2019), proteomics/protein abundances (Aslam et al., 2017; Gorgoni et al., 2016), and smFISH (Pichon et al., 2018; Cui et al., 2016) provide snapshot bulk data in high quantity, at the detriment of obscuring the temporal dynamics of individual mRNA molecules. Pulsed SILAC, PUNch-P, BONCAT/QuaNCAT allowed mass spectrometry quantification of recently translated protein abundances via treatment with noncanonical amino acids (Howden et al., 2013; Eichelbaum et al., 2012). Methods such as FUNCAT/SUnSET bridged the gap of detecting active mRNA translation as well as subcellular location by labeling nascent peptide chains and imaging in fixed cells (Dieck et al., 2012; David et al., 2012). FUNCAT-PLA and Puro-PLA then provided spatial resolution and imaging of the recent protein production via detection with a proximity ligation assay (PLA) (Dieck et al., 2015). Despite their innovations, these techniques require fixation or lysis of the cells of interests. As a consequence, none of these methods are able to capture long-term temporal imaging of translation at a single-molecule level of the same mRNA molecules (Chekulaeva and Landthaler, 2016).

Since 2016, Nascent Chain Tracking (NCT) has provided experimentalists with a technique to study single, actively translating mRNA transcripts and quantify their dynamics with the use of a dual fluorescent labeling system (Cialek et al., 2020; Pichon et al., 2018; Morisaki et al., 2016; Wu et al., 2016; Yan et al., 2016; Pichon et al., 2016). The key to this technique is multiple epitope sequences that are placed on a tag within the coding region of the studied mRNA; After this tag region is translated, fluorophore-conjugated intrabodies bind to the growing nascent polypeptide chain resulting in an amplified fluorescent spot. After translation is completed, fluorescently-tagged single proteins are free to diffuse away. The mRNA molecule is tagged in the 3’ untranslated region with a hairpin loop repeat system recognized by fluorophore-conjugated MS2 or PP7 coat proteins, conferring a constant intensity in a separate color channel for tracking purposes. The combination of these two elements gives a co-localized, diffraction-limited, two-color spot trajectory denoting an mRNA location and a live nascent chain activity readout. NCT has been utilized to investigate many processes of interest such as mRNA frameshifting (Lyon et al., 2019), mRNA IRES-mediated translation (Koch et al., 2020), mRNA decay (Horvathova et al., 2017; Hoek et al., 2019), translation suppression during cellular stress (Moon et al., 2019), and mRNA spot to spot heterogeneity (Boersma et al., 2019). NCT has also proven beneficial to extract important biophysical parameters, including elongation rates, initiation rates and ribosomal densities (Morisaki et al., 2016; Wu et al., 2016; Yan et al., 2016; Pichon et al., 2016), and microRNA mediated decay (Kobayashi and Singer, 2022; Cialek et al., 2022).

Notwithstanding NCT’s current adoption and importance, application of NCT to understand how different cellular signals affect translation of different mRNA is currently prevented by strong limits on the fluorophore color palette. Currently, only three or four resolvable colors exist in most microscope settings; one color is dedicated to tracking the mRNA, leaving only two or three resolvable colors to design experimental setups and constructs. Use of too many fluorescent probes with similar emission spectra leads to imaging issues such as light bleed through into each channel. Additional laser wavelengths also limits the frame rates that one can utilize due to the time required to switch laser or filters between each frame. While this has not proven detrimental to previous NCT experiments that have explored one (Morisaki et al., 2016; Yan et al., 2016; Wu et al., 2016) or two (Lyon et al., 2019; Boersma et al., 2019; Koch et al., 2020) translation products at a time, there is an anticipation that this limitation will become a roadblock for designing future, more elaborate experiments, particularly for processes involving differential control of multiple mRNAs under cellular stimuli. Specifically, we speculate that the current form of NCT technology has already reached its peak in experiments where two different genes are correctly detected and differentiated on the same cell (Lyon et al., 2019; Koch et al., 2020; Boersma et al., 2019).

Recently, machine learning has enjoyed an explosion of applications in biological and biomedical imaging contexts (Tchapga et al., 2021; Haque and Neubert, 2020). Convolutional neural networks have achieved the state-of-the-art performance on a wide variety of classification tasks such as speech recognition (Palaz et al., 2015), computer vision (Jarrett et al., 2009), natural language processing (Liao et al., 2017), and in myriad biomedical contexts (Zhu et al., 2018; Feeny et al., 2020). For the purpose of our work, we utilize 1D convolutional neural networks to classify signals from two mRNA species recaptured from a realistic noise model and realistic mRNA translation model. One dimensional convolutional neural networks (CNNs) are well equipped to handle 1D signals for classification and see frequent use for applications across the biomedical field such as ECG (Murat et al., 2020) and EEG signals (Xie and Oniga, 2020).

Applying machine learning to NCT experiments cannot be done outright as of time of writing due to a lack of large and standardized data sets, and it is not clear exactly how much data, and under which conditions, would be needed to build successful classifiers. To more efficiently explore these questions, we generate large sets of realistic NCT experiment simulations using a new pipeline, as detailed below. In brief, translation of nascent proteins and their corresponding Fluorescent intensity signals are modeled with a codon-dependent Totally Asymmetric Simple Exclusion Process (TASEP) to consider elongation rate changes due to codon selection along each mRNA transcript, as well as ribosomal collisions. In this paper, we use our previously described mRNA translation mechanistic model–a full comprehensive explanation of this model and its parametrization can be found in (Aguilera et al., 2019). Fluorescent intensity signals from the mechanistic model are then combined with our realistic video rSNAPed pipeline, which applies a point spread function and adds a simulated cell background with microscope noise calibrated from real videos. mRNA molecules freely diffuse with a set diffusion rate within the simulated cell mask to simulate Brownian motion. The resulting intensity from the simulated videos are then processed as if they were real images to generate “simulated nascent chain tracking” data. Through this means, we construct a controllable “experiment” whose measurement is a corrupted fluorescent signal, where we can easily control various factors such as signal-to-noise ratio (SNR), spot size, diffusion rates and mechanisms, mRNA initiation and elongation rates, and imaging conditions (frame rate / frame interval and number of frames taken).

In this paper, we use these simulations to demonstrate that high classification accuracy is achievable in principle using a machine learning approach across a large range of biophysical and experimental parameters, even if two mRNAs are utilizing identical tagging approaches and fluorescent colors. To extend NCT color palette, our computational pipeline of mRNA translation modeling, spot simulation, spot tracking, and machine learning uses as features various statistics of the spot’s intensity fluctuations, such as their moments, relative intensity ratios, and decorrelation times to discriminate between different mRNA species. This type of “temporal multiplexing” could radically expand the number of mRNAs imaged in a cell. Different color fluorophores could be held in reserve for mRNAs whose characteristics are too similar to each other, or eschewed altogether to increase microscope imaging speed and decrease experiment cost. The entire pipeline can be used to explore potential NCT experimental designs without using valuable lab time and resources. We envision that the proposed model-based strategies to tag multiple mRNAs using single color tags will add new possibilities for future experimental NCT investigations.

## 2 Methods

### 2.1 Simulated NCT experiment pipeline

Figure 1 graphically describes the current study’s computational pipeline, which combines the RNA Sequence to NAscent Protein SIMulator (rSNAPsim) and rSNAP Experiment Designer (rSNAPed) scientific libraries to generate synthetic training and testing data sets for multiple experimental conditions. rSNAPsim is a Python module that provides simulated fluorescent intensity traces from a given mRNA transcript using a codon-dependent TASEP to simulate ribosomal elongation (Aguilera et al., 2019). This mechanistic model for translation assumes two parameters: the ribosomal *initiation rate*, *k*_i_, is defined as the average number of ribosomes to initiate translation per second, assuming that other ribosomes are not blocking the initiation site. The *elongation rate*, *k*_e_ is defined as the global stepping rate (aa/s) averaged over all coding regions in the genome, again assuming no ribosome-to-ribosome exclusion. Because the actual local stepping rates depend the specific codon usage of an mRNA, and because rSNAPsim includes a 9-codon ribosome exclusion footprint that prevents two ribosomes from occupying the same site at the same time, the effective stepping rate for each mRNA is based on the specific sequence of the mRNA. For full details on the rSNAPsim model, including calculations for the individual codon elongation rates, see (Aguilera et al., 2019). We have shown previously that the rSNAPsim module can reproduce much of the fluctuation statistics observed in NCT translation experiments (Aguilera et al., 2019; Lyon et al., 2019; Koch et al., 2020). However, the movement of NCT spots across the cell background can lead to large drops in intensity when, for example, a spot leaves a bright nuclear region and enters a dimmer cytoplasmic region. These movements from areas of high to low or low to high backgrounds cause intensity fluctuations that are not due to the translation process itself but are, nevertheless, always present in real experimental data.

**Figure 1:**
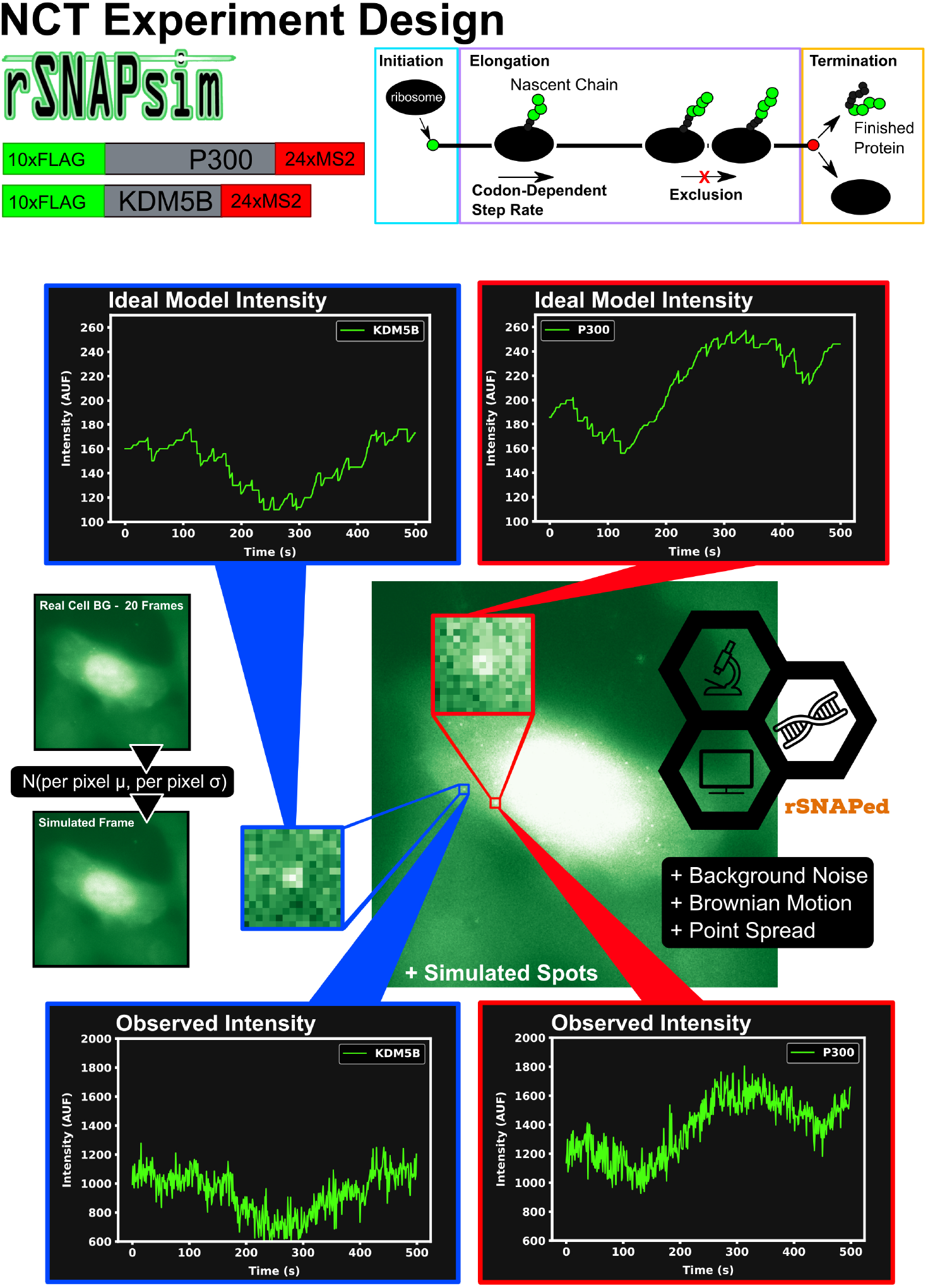
Overview of approach to simulate Nascent Chain Tracking data and assign labels. rSNAPsim provides simulated NCT fluorescent intensity trajectories from a codon-dependent TASEP model for each mRNA spot. rSNAPed adds experimental spatial movement (Brownian motion) and temporal noise by introducing a point spread function for each spot. Simulated cell background frames are generated randomly from a per pixel Gaussian distribution with their means and standard deviations taken from 20 frames of real blank cell backgrounds. Spots in videos are processed with the disk and doughnut method to generate simulated NCT intensity data.

rSNAPed is a second python module that accounts for artifacts of microscopy, motion relative to a heterogeneous cellular background, and image processing effects. rSNAPed combines the rSNAPsim model intensity prediction with a point spread function, a controllable signal-to-noise ratio, simulated cellular background based on experimental video of non-labeled cells, and simulated motion for the NCT spots. To create a video for any length of time, rSNAPed takes 20-frame videos from one of seven unlabeled cells, randomly rotates and flips the videos and uses pixel-by-pixel statistics to formulate distributions from which to generate new simulated frames. Specifically, for each simulated frame, rSNAPed uses each pixel’s empirical mean and standard deviation to draw a new Gaussian distributed value for the respective pixel^1^ Supplemental Figure S1 provides a comparison of real cell background video and a simulated video from rSNAPed. Simulated mRNA spots are added to this cell background by simulating a point spread function on a 3×3 pixel patch, which is centered at a position that moves according to Brownian motion. After simulating the NCT experiment, videos are processed to find intensity trajectories using the “disk and doughnut” method (Lyon et al., 2019), where the instantaneous signal is quantified as the difference between the average of the disk (3×3 patch centered at the spot) compared to the average of the doughnut (9×9 patch excluding the 3×3 disk patch). Units of intensity are reported as “units of mature protein” or UMP, which is calculated as the number of complete epitope tags in the NCT spot (i.e., if a simulation has two ribosomes downstream of a 10xFLAG tag and one halfway through the tag, the intensity at that time is 25 epitopes or 2.5 UMP).

The combination of rSNAPsim and rSNAPed allows generation of vast amounts of synthetic video in different situations (e.g., for different mRNA sequences, different biophysical parameters, and different imaging conditions) that can match the translation statistics and spatial heterogeneity that an NCT spot would experience as it moves around a cell. For each mRNA in each condition, we generate 2500 simulated NCT trajectories (5000 total trajectories for two mRNA types in the same video). Theses trajectories are collated from 100 independent NCT simulated cells, each containing 25 spots of each mRNA for 3000 seconds at one second resolution (25 spots × 2 classes × 100 cells = 5000 total NCT spots for 3000 seconds). Smaller data sets can be generated as needed from these full length data sets by slicing the 1 second × 3000 frame video trajectories to the desired frame interval and number of frames.

By generating such data for multiple different potential experimental conditions, we can train and test our machine learning methods to ascertain which feasible experimental conditions are most favorable to allow for successful classification.

### 2.2 Machine Learning

Figure 2A shows the machine learning architecture used to classify mRNA spots. Given two different mRNA species in the same cell and NCT observations that rely on identical tags, one could attempt to classify the mRNA based on their intensity signals, their particle sizes, or their x, y, and z coordinates over time (with z having poorer resolution than x and y). Of these, we focus on intensity signals, which contain both statistical moments, such as the signal means and variances, as well as signal frequency content, similar to that which could be obtained with methods like spectrogram analysis or fluctuation correlation spectroscopy (FCS). We apply convolutional neural networks to classify the NCT simulation data based on one or both types of signal intensity inputs. First, to prioritize extraction of features related to intensity statistics, we use min-max normalization of all intensity signals in the same video. Let *I_i_*(*t*) denote the intensity for spot *i* ∈ (1, 2,…) and at frame number *t* ∈ [0, 1*,…, T* − 1] that has been collected from a given NCT experiment or simulation video. Define *I*_max_ and *I*_min_ as the maximum and minimum spot intensities across all spots and all times in the video:

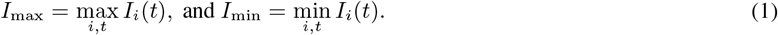

**Figure 2:**
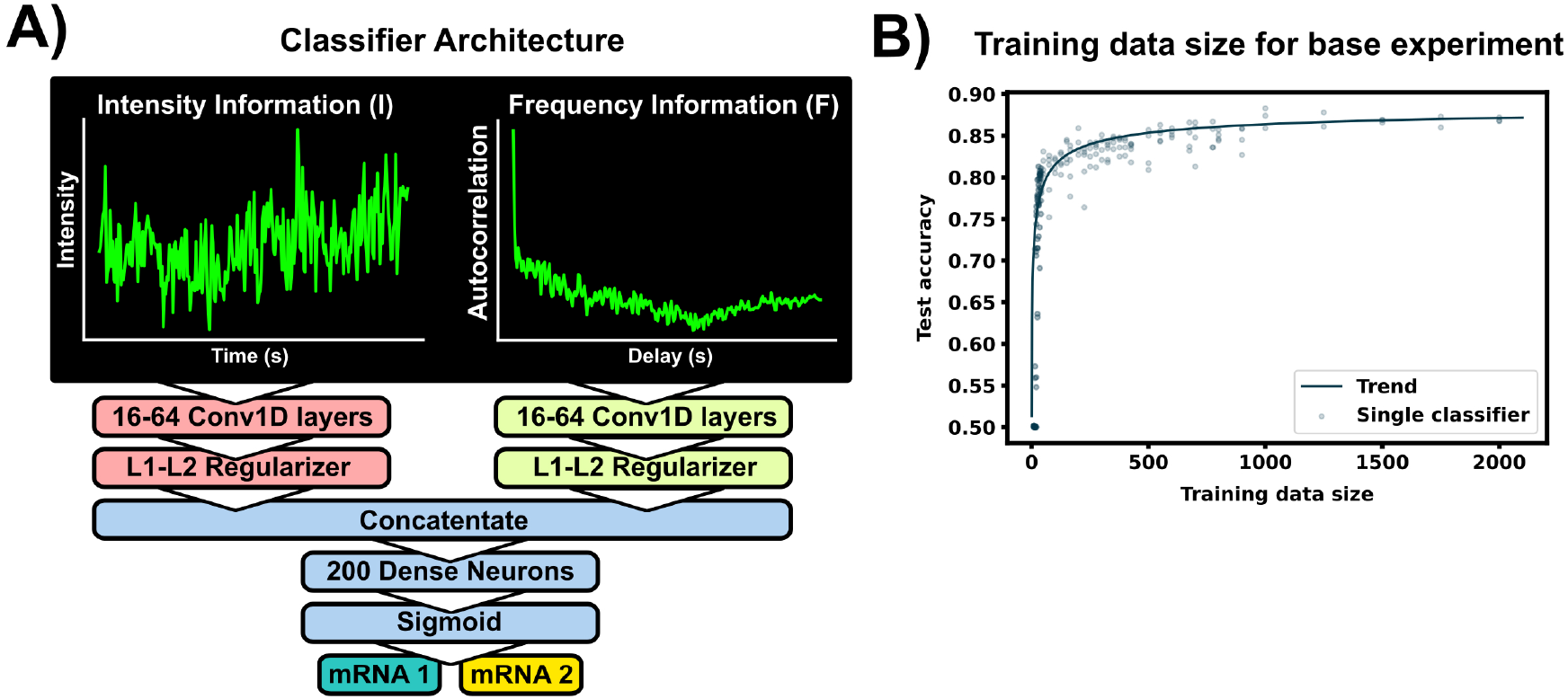
Machine learning to classifier mRNA spots. (A) The ML model consists of two separate convolutional layers – one receiving a normalized fluorescent trajectory and the other an intensity autocorrelation. The filter outputs are regularized and concatenated for a dense layer of 200 neurons for classification. Fundamentally, this architecture learns off frequency and intensity information from the NCT trajectory. B) Accuracy of the architecture for different training data sets. A total of 4000 unique spot trajectories were split into 2-10 independent training data sets of the specified size. A classifier was trained with each training data set and tested on the same withheld validation set of 1000 NCT spots. A trend-line was added by fitting a Hill function to the test accuracy average across 15 bins in training data size 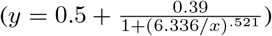. The architecture was applied on simulated P300 and KDM5B trajectories from the selected base experimental condition (5s frame interval, 64 frames, 0.06 1/s initiation rate, 5.33 aa/s elongation rate).

The min-max normalization of each signal, *I_i,_*_norm_, which scales features from zero to one for later convenience in machine learning, is computed as:

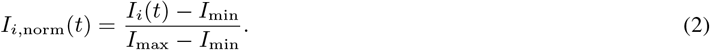

Second, to prioritize features related to fluctuation frequencies, we use the normalized empirical auto-correlation function for each spot, *G_i_*(*τ*), which is defined as the sample covariance between the signal fluctuation at frames *t* and *t* + *τ*, is calculated as:

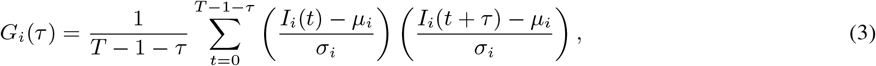

where *t* is an index corresponding to the frame number, *τ* is the correlation lag time; *T* is the total number of frames; and *μ_i_* and *σ_i_* are the i^th^ signal’s trajectory average and standard deviation, respectively (Morisaki et al., 2016; Coulon and Larson, 2016).

The normalized inputs from Equations 2 and 3 are each passed to their own separate convolutional 1D layer and subsequent max pooling layer for feature extraction. For convenience, the two convolutional layers have the same size filter kernels and amount of filters. The extracted feature vectors from each input are concatenated and passed into one fully connected layer with a cross-entropy objective to classify the mRNA. The entire network (2 conv1D, 2 maxpooling, 1 dense) was trained end to end, and elastic net regularization is used to reduce over-fitting (Zou and Hastie, 2005). During training, data was split with an 80:20 ratio into training and testing sets, and training was performed with a 3-fold cross validation using a random search to select hyperparameters. Specifically, we searched over the possible combinations of options listed in Table 1 (Bergstra and Bengio, 2012), and selected the hyperparameter set that maximized validation accuracy. After training and hyperparameter selection, the final architecture was tested on the 20% withheld test data to quantify the model performance on unseen data.

**Table 1:** Hyperparameter optimization grid

| Hyperparameter | Variable range |
| --- | --- |
| kernel size | 1x3, 1x5, 1x7 |
| number of kernels | 16, 32, 64 |
| batch size | 16, 32, 64 |
| epochs | 50, 100 |

### 2.3 Hardware

All machine learning experiments were done with TensorFlow 2.2.0 on 2x NVIDIA Geforce 2080 Supers. Simulated NCT experiments were generated on an AMD Ryzen Threadripper 3970X 32-Core Processor utilizing 8 threads for each simulated NCT experiment generation.

Time for generating one 5000 spot NCT experiment with base experimental conditions: ≈ 2h 44m

Time for generating one 50-spot cell with base experimental conditions (short output + not saving video): ≈ 197 seconds

### 2.4 Microscopy

The seven background videos used for video generation in the rSNAPed were captured using a custom-built wide-field fluorescence microscope with a highly inclined illumination regime with 488, 561, and 637 nm excitation beams (Vortran) ((Tokunaga et al., 2008; Morisaki et al., 2016)). An objective lens of 60X, NA 1.49 oil immersion, Olympus, was used. The emission signals were split by an imaging grade, ultra-flat dichroic mirror (T660lpxr, Chroma) and detected using two separate EM-CCD (iXon Ultra 888, Andor) cameras via focusing with 300 mm tube lenses ultimately producing 100X images with a 130 nm/pixel resolution. JF646 signals were detected with the 637 nm lasers and the 731/137 nm emission filter (FF01-731/137, Semrock). Cy3 signals were detected with the 561 nm lasers and the 593/46 nm emission filter (FF01-593/46, Semrock)

The lasers, the cameras, and the piezoelectric stage were synchronized via an Arduino Mega board (Arduino) and image acquisition was done with open source Micro-Manager software (Edelstein et al., 2014). An imaging size of 512×512 pixels^2^ was used and exposure time was set to 53.63 msec. This resulted an imaging rate of 13 Hz with 23.36 msec readout time and 13 Z-stacks were captured at 500nm step size.

U-2 OS cells were loaded with Cy3-FLAG Fab and JF646-Halo-MCP 6 to 10 hours prior to imaging. Right before imaging, cells were transferred into the stage top incubator set to a temperature of 37°C and supplemented with 5% CO_2_ (Okolab). Acquired 4D (xyzt) images were processed to 3D (xyt) maximum intensity projections for the rSNAPed image generation.

## 3 Results

To begin our exploration of the potential to use NCT signal intensity fluctuations to differentiate mRNA species, we choose baseline experiment using P300 (7257 NT) and KDM5B (4647 NT) mRNA, each with a 10xFLAG epitope tag. Both constructs are assumed to have equal initiation and elongation rates of *k_i_* =0.06 1/s and *k_e_* =5.33 aa/s, and both are assumed to be images for 64 frames with a rate of one frame every five seconds.

### For typical mRNA and experiment designs, large training data sets may be needed to build an accurate classifier

To see how much training data are needed to build an accurate classifier, we use the baseline mRNA designs and experimental conditions (with 64 frames), and we trained our architecture with progressively increasing training data sizes. Figure 2B shows the resulting accuracy on a withheld validation set of 1000 NCT spots, when the model is trained on independent training sets of the different sizes. The accuracy of the classifier levels off at about 87% when the training data set reaches 800-1000 NCT spots. Unfortunately, such a large amount of data is approaching unfeasible in current NCT laboratory settings, highlighting the need for using mechanistic simulation to supplement data when training a classifier. It is important to note that this result is for the experimental base conditions listed in Table 2; as we will discuss below, other parameter combinations can increase or decrease classification difficulty and result in variations for the amount of training information required.

**Table 2:** Selected base experimental conditions, statistics are calculated before microscope noise addition via rSNAPed.

|  | Gene 1 - KDM5B | Gene 2 - P300 | Imaging conditions |
| --- | --- | --- | --- |
| **Parameters** | $L$: 4647 NT<br>$L_{tag}$: 1011 NT<br>$k_e$: 5.333 aa/s<br>$k_i$: 0.06 1/s | $L$: 7257 NT<br>$L_{tag}$: 1011 NT<br>$k_e$: 5.333 aa/s<br>$k_i$: 0.06 1/s | 64 frames<br>5s frame interval<br>KDM5B SNR: 6.2<br>P300 SNR: 8.9 |
| **Statistics** | $\mu_I$: 19.3 UMP<br>$\sigma_I^2$: 18.7 UMP<br>$t_{dwell}$: 354 s | $\mu_I$: 28.2 UMP<br>$\sigma_I^2$: 28.5 UMP<br>$t_{dwell}$: 517 s | |

\* NT = nucleotides, \* aa = amino acids

### Realistic simulations of nascent chain tracking for KDM5b and P300 mRNA reveal that two mRNA can be distinguished using only their fluorescence intensity fluctuations

As a proof of concept and to further explain the process of labeling spots by their behaviors rather than by their colors, Figure 3 presents a example of our machine learning pipeline. Two simulated cells (Figure 3A, top) are generated using the baseline conditions but with double the amount frames (128 instead of 64, but with the same 5s interval between frames) to better highlight the ribosomal dwell-time difference between the two mRNAs. The two cells can be processed with a disk and doughnut approach (see Methods) to extract fluorescence intensity trajectories for each of the 100 spots (Figure 3A, bottom).

**Figure 3:**
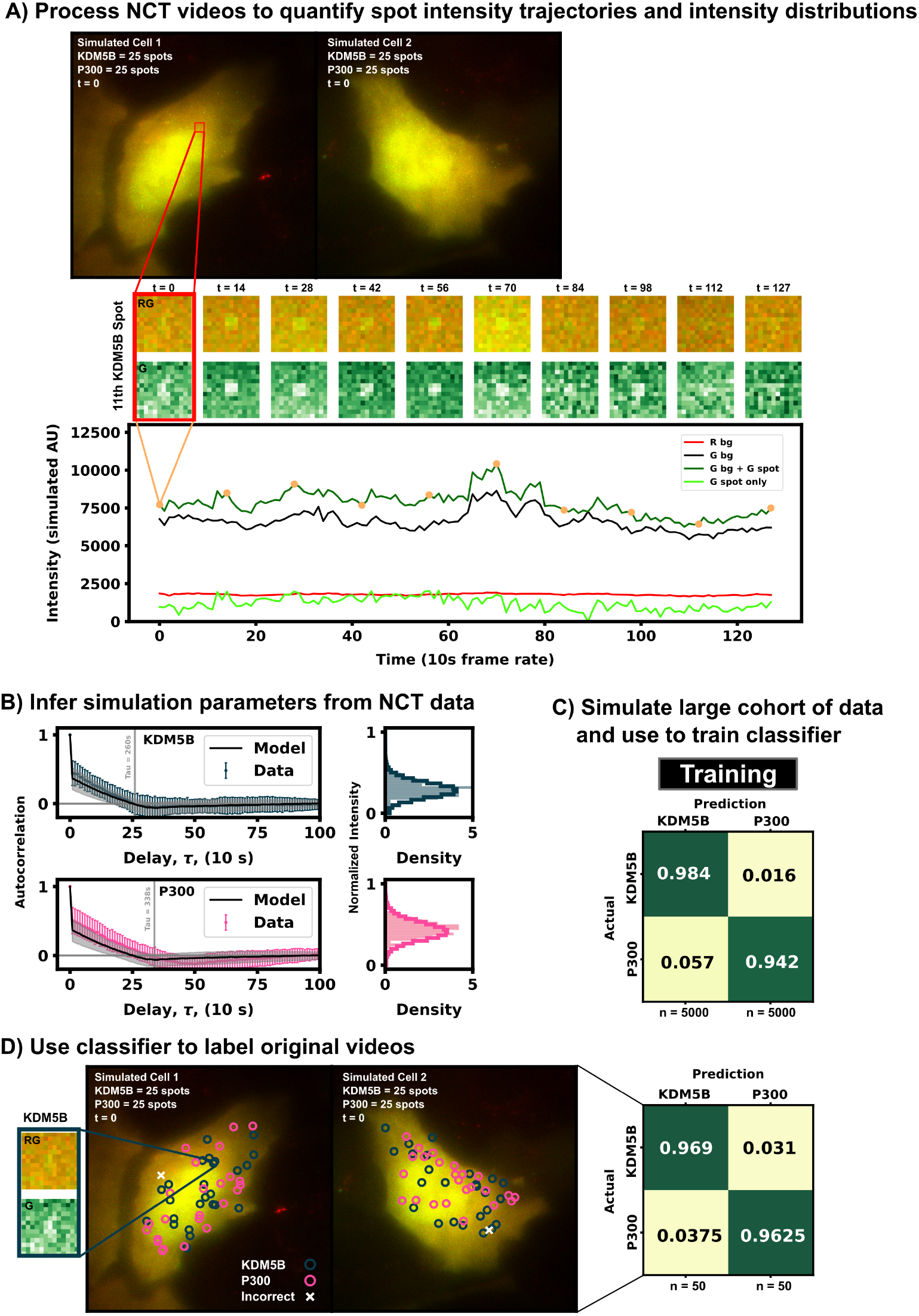
Example for labeling identically-tagged mRNAs in a simulated NCT experiment. (A, top) Two simulated cells with 25 KDM5B and 25 P300 spots translating at identical biophysical parameters. (a, bottom) Spot 11 in Cell 1 is highlighted in the intensity trace, showing the red background, green background, and extracted spot intensity via the disk and doughnut method. (B) Model parameters (*k*_i_ and *k*_e_) are inferred by fitting auto-correlation functions and intensity distributions. (C) A classifier is trained with a large cohort of simulated data generated with the inferred parameters. (D) This classifier can then be used to label the original data or any subsequent data taken.

In practice, biophysical parameters could be estimated from these NCT measurements of fluorescence intensity trajectories (i.e., from the intensity distributions and autocorrelations) as done previously in (Aguilera et al., 2019) and illustrated in Figure 3A. For simplicity of description, we assume that these parameters are known, although we will relax this assumption later in the investigation. Using these assumed parameters, we use our computational pipeline to simulate a training data set of 5000 NCT spots from simulated NCT experiments, and we use this simulated data to train a classifier (Figure 3C). With this classifier, the user can then finally label their original data artificially or use the trained classifier to label any newly collected data (Figure 3D).

### Simulations can reveal which aspects of experimental data are most informative for multiplexed mRNA classification

For two different NCT spots within the same cell and under the same experimental conditions, our dual input architecture (Figure 2A and Methods) utilizes both relative intensity differences and signal frequency content to classify spots. Using both features allows for improved classification robustness across parameter space. There are experimental conditions and biophysical parameters where intensity information is more useful, conditions where frequency is more informative, and conditions where a mixture of both information sources is utilized by our architecture. To highlight this, we simulated three separate data sets: one with markedly different intensity distributions, one with mostly overlapping distributions, and one with nearly identical intensity distributions, Table 3. For each of of these conditions (which have different translation initiation rates), both P300 and KDM5b use the same average elongation rate, but because the genes have different lengths and different codon usages, they exhibit different ribosomal dwell time.

**Table 3:** Long Imaging Time Datasets

| Comparison Analysis | Variable Range | Constants |
| --- | --- | --- |
| **Imaging Dataset - Identical $I_\mu$**<br>$I_{\mu,both} \sim 4.7 \text{ UMP}$ | $k_{i,KDM5B}: 0.014139 \text{ s}^{-1}$<br>$k_{i,P300}: 0.009676 \text{ s}^{-1}$ | * Gene 1: P300 (7257 NT)<br>* Gene 2: KDM5B (4647 NT)<br>* Frame interval: 1 every 1 seconds<br>* Frames: 24000<br>* $k_e: 5.33 \text{ aa} \cdot \text{s}^{-1}$<br>* D: 0.21 pixels<sup>2</sup>/s<br>* SNR: 3 |
| **Imaging Dataset - Similar $I_\mu$**<br>$I_{\mu,KDM5B} \sim 6.0 \text{ UMP}$<br>$I_{\mu,P300} \sim 4.7 \text{ UMP}$ | $k_{i,KDM5B}: 0.01860 \text{ s}^{-1}$<br>$k_{i,P300}: 0.009676 \text{ s}^{-1}$ | * Gene 1: P300 (7257 NT)<br>* Gene 2: KDM5B (4647 NT)<br>* Frame interval: 1 every 1 seconds<br>* Frames: 24000<br>* $k_e: 5.33 \text{ aa} \cdot \text{s}^{-1}$<br>* D: 0.21 pixels<sup>2</sup>/s<br>* SNR: KDM5B (3.9) and P300 (3) |
| **Imaging Dataset - Different $I_\mu$**<br>$I_{\mu,KDM5B} \sim 13.7 \text{ UMP}$<br>$I_{\mu,P300} \sim 4.7 \text{ UMP}$ | $k_{i,KDM5B}: 0.04242 \text{ s}^{-1}$<br>$k_{i,P300}: 0.009676 \text{ s}^{-1}$ | * Gene 1: P300 (7257 NT)<br>* Gene 2: KDM5B (4647 NT)<br>* Frame interval: 1 every 1 seconds<br>* Frames: 12000<br>* $k_e: 5.33 \text{ aa} \cdot \text{s}^{-1}$<br>* D: 0.21 pixels<sup>2</sup>/s<br>* SNR: KDM5B (8.9) and P300 (3) |

We applied our architecture to each of these data sets across a large swath of imaging conditions (i.e., different numbers of frames and frame intervals) to show our architecture’s ability to self select which features to use for classification as well as highlight which imaging conditions are ideal for these experiments conditions. Figure 4A (left) shows that the first condition (different dwell times and different intensity distributions) is trivial to classify – Simply looking at which spots are brightest and which are dimmest is sufficient to reach than 90% accuracy in just a few frames. Figure 4A left) also shows that for mRNAs with such different intensity distributions, fluctuation frequencies are less informative, and the optimal frame interval should have as long a delay as possible so that each measurement is as statistically independent as possible.

**Figure 4:**
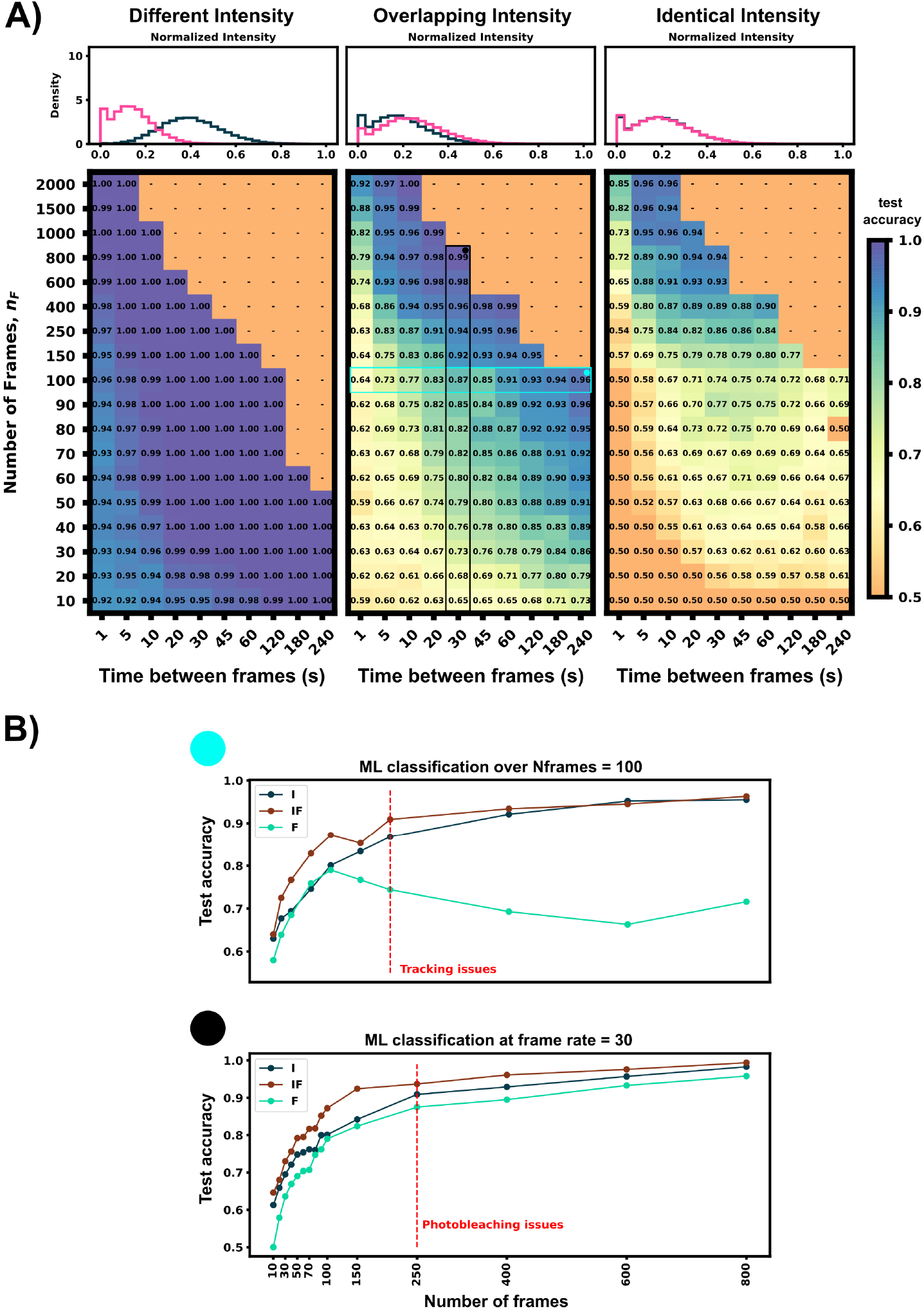
Classification accuracy versus imaging conditions and differences in mRNA intensity mean. (A) Accuracy versus frame interval and number of frames. (Left) For constructs with substantially different intensities, the classifier requires only a few frames for a high classification accuracy. (Middle) Overlapping, but non-identical, intensity conditions leverage both frequency and intensity information for classification. (Right) Identical intensity conditions can only classify using frequency information, which requires an ideal frame interval. (B) Accuracy of ML using Intensity only (I), Frequency only (F), and both (IF) versus frame interval (top) or number of frames (bottom). Plots for (IF) correspond to the vertical and horizontal regions highlighted in Panel A, middle.

Conversely, identical intensity conditions can be obtained by tuning the initiation rates such that each mRNA has the same ribosomal occupation average over time, and thus almost identical intensities (there is a slight difference due to codon dependence and ribosomal occupation probabilities per codon leading to varied occupation downstream of the tag region across each mRNA’s length). In this condition, our classifier can only learn on the autocorrelation function and frequency information. Expected ribosome dwell times for P300 and KDM5B under these conditions are approximately 517 seconds and 353 seconds, respectively – a statistic that is available to the classifier through the autocorrelation of its intensity fluctuation. Figure 4A (right) shows the test set classification accuracy as a function of video frame intervals and total number of frames. This plot highlights a clear region of image settings that would be sufficient to capture enough frequency information for classification, ≈20-30 seconds between frames for over 150 total frames. To be effective, the frequency-based classifier needs enough frames at the right intervals to sample the auto-correlation of ribosomal movements. If the total video time is too short, too few ribosomes will complete translation, and the NCT intensity signal would remain almost fully correlated with itself. As a result, one would have insufficient number of independent data points with which to calculate an effective autocorrelation. Conversely, if one observes the process too slowly (i.e., approaching or exceeding ribosomal dwell times), each frame would sample an independent set of ribosomes from the previous frame and any observed correlations would arise only from artifacts of imaging noise.

Figure 4A (middle) shows an experimental condition where intensity distributions are similar enough such that both frequency and intensity information provide useful information for classification. The ideal imaging still uses an intermediate frame interval needed to capture the frequency differences, but classification is bolstered by the intensity information across the whole parameter space, with 10 frames at any frame interval being sufficient to provide 60% accuracy.

To further probe how intensity distributions and frequency information each contribute to ML classification, Figure 4B shows each half of the architecture applied separately to 100 frames at varying frame intervals and varying amounts of frames at a 30 second frame interval (Highlighted row and column of Figure 4A, middle panel). Both individual halves Intensity distribution, (I), and Frequency, (F), have roughly a similar accuracy until frame interval grows too large to obtain a good sampling of the autocorrelation function (60s). When both features are available, (IF), the classifier has a marked improvement in accuracy compared to the individual halves. It is important to note that we selected these specific experimental conditions such that there is partial information in both the autocorrelation and the intensity moments. If the experiment is designed such that either frequency or intensity is more informative, the proposed ML architecture adapts to rely more on that type of information and including the other will have marginal or no effect on validation accuracy (e.g., Figure 4A left or right panel).

Although classification requires collection of a sufficient video length and at high enough temporal resolution, other experimental considerations are missing in this first analysis but must be taken into account to constrain imaging conditions in a real laboratory setting. Taking too many frames or a too short a frame interval may increase photobleaching effects that will require re-calibration or computational correction or may necessitate the use of a lower laser power, which will reduce signal strength. Conversely, choosing too low a frame interval (i.e., longer delays between images) could lead to spot tracking issues if particles diffuse too fast in comparison to the frame rate or if there is too high a density of overlapping particles. This trade-off between too fast and too slow a frame interval can be partially ameliorated by tracking only on the RNA tag channel with a higher frame interval and imaging in the protein channel with a slower rate, but this solution requires more complex steps for image collection and processing.

### Simulated data is ideal for testing different experimental and biological conditions to probe the possibilities and limitations of NCT multiplexing

Now that we have a computational pipeline to generate simulated data and a flexible classifier to train using both intensity and frequency information, we can explore multiple parameter spaces to guide experimental design toward conditions that are more conducive for accurate classification. Additionally, by generating data over a large parameter swath, we can examine multiple experiment setups that are unresolvable to obtain insight for what mitigation strategies an experimentalist could take to improve classification results.

In this study, we limit the scope of exploration to five key variables (Table 4): mRNA length (*L_mRNA_*), frame interval (time in seconds between frames, *FI*), number of frames used (*n_F_*), ribosomal initiation rate (*k*_i_) and ribosomal elongation rate (*k*_e_). We choose these specific parameters because they are the most experimentally relevant as they influence how long of a video to take and what types of mRNA constructs to design. Additionally, these five parameters directly influence one or both of the two statistics we are using as learning features, specifically the decorrelation times and intensity levels. The selected parameters to explore and a list of dynamics affected by each of these parameters are provided in Table 4. For example, an increase in mRNA length has a corresponding increase in ribosomal dwell time and therefore fluorescent intensity of a mRNA spot.

**Table 4:**
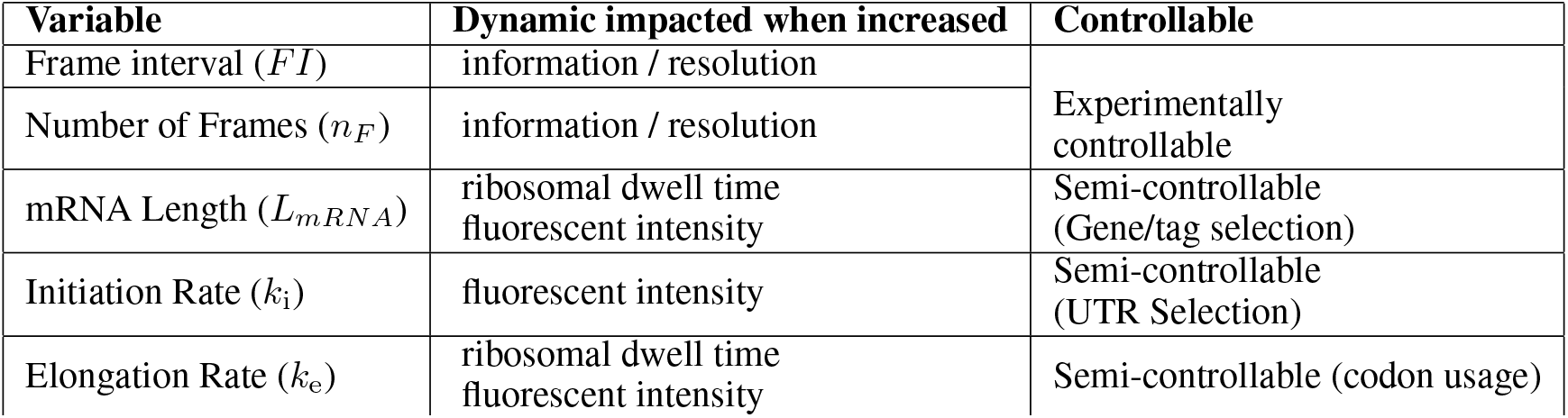
Dynamics affected by selected variables to investigate with the NCT ML pipeline

| Variable | Dynamic impacted when increased | Controllable |
| --- | --- | --- |
| Frame interval ($FI$) | information / resolution | Experimentally controllable |
| Number of Frames ($n_F$) | information / resolution | |
| mRNA Length ($L_{mRNA}$) | ribosomal dwell time<br>fluorescent intensity | Semi-controllable (Gene/tag selection) |
| Initiation Rate ($k_i$) | fluorescent intensity | Semi-controllable (UTR Selection) |
| Elongation Rate ($k_e$) | ribosomal dwell time<br>fluorescent intensity | Semi-controllable (codon usage) |

Although for the sake of brevity, we focus on the five parameters presented in Table 4, we note that our proposed simulation and classification pipeline is general and can be used to explore many other mechanisms of the translation or the imaging system. Other parameters of potential interest may improve or worsen classification accuracy. These include effects such as imperfect spot detection and tracking, non-equilibrium dynamics (all mRNA simulations in this paper are at steady state), differing diffusion dynamics (spatial considerations from cell morphology or dependence on mRNA translation state), or photobleaching effects. For example, to focus our analysis on our current parameters of interest, all data are recaptured using the specific coordinates of the simulated spots within the rSNAPed module; i.e., we are not relying on a spot detection and tracking algorithm. However, rSNAPed also includes a tracking algorithm (Allan et al., 2021) that one could use to explore the effects that imperfect tracking has on the generated videos. Similarly, all translation models are run until they reach steady state (burn in time: 1000 seconds) before they are used to generate NCT spots, and photobleaching is not added to the videos, but rSNAPed allows for simulation of non-stationary conditions (e.g., Harringtonin treatment, or FRAP) or the inclusion of photobleaching or other effects on the signal to noise ratio if required. Finally, all spot motion is simulated using normal Brownian motion with a constant diffusion rate of 0.925 *μm*^2^/s (0.55 pixel^2^/s) but settings for rSNAPed can easily be changed to allow for anomalous or temporal or mRNA state-dependent diffusion rates. In addition, beyond using just intensity and frequency, one could potentially improve classification using additional features for machine learning such as x, y, and z spatial positions or spot velocities.

Figure5(top) summarizes the set up for four parameter sweeps designed to explore the effects of selected parameters on ML classification: (CT1) compares pairs of mRNA with many different lengths, (CT2) compares two mRNA with many different combinations of shared elongation and initiation rates, (CT3) compares two mRNA each with different elongation rates, and (CT4) compares two mRNA each with different initiation rates. For each comparison, mRNA translation simulations use parameters selected from a survey of literature reporting experimentally measured rates (Morisaki et al., 2016; Wang et al., 2016; Wu et al., 2016; Pichon et al., 2016; Yan et al., 2016), and all parameters are shown in Table 5. For CT1, the mRNA length range selected (1200 - 7257 nt) covers 53% of the lengths in the human consensus coding sequences (current CCDS nucleotide release - 11.28.2021 (Pujar et al., 2018)). A baseline NCT experiment was selected as 10xFLAG-p300 vs 10xFLAG-KDM5B with an initiation rate and elongation rate of 0.06 1/s and 5.33 aa/s at imaging conditions of 64 frames with a 5s frame interval. When parameters are held constant in the comparison analysis they are held to these values. The results of the various comparison tests are shown in Figures 5A-D and 6A-D, to be discussed individually below, and all full data sets are provided at: https://www.dropbox.com/scl/fo/cj4fd2o0nr5dfstngyh83/h?dl=0&rlkey=363ka3o1ur8jvywhh5mvolxqg.

**Table 5:** Comparison analyses parameters

| Comparison Analysis | Variable Range | Constants |
| --- | --- | --- |
| **mRNA Length vs mRNA Length** | mRNA length (without tag):<br>1200 NT - 7257 NT<br>RRAGC - 1200<br>ORC2 - 1734<br>LONRF2 - 2265<br>EDEM3 - 2799<br>TRIM33 - 3333<br>MAP3K6 - 3867<br>COL3A1 - 4401<br>KDM5B - 4657<br>KDM6B - 4932<br>PHIP - 5466<br>DOCK8 - 6000<br>P300 - 7257 | * Frame interval: 5s<br>* Frames: 64<br>* $k_i$: $0.06 s^{-1}$<br>* $k_e$: $5.33 aa \cdot s^{-1}$<br>D: $0.21 pixels^2/s$<br>SNR: 3.7 (RRAGC) - 17.6 (P300) |
| **$k_e$ (both mRNAs) vs $k_i$ (both mRNAs)** | $k_i$: $0.01 - 0.1 s^{-1}$<br>$k_e$: $2 - 12 aa \cdot s^{-1}$ | * Gene 1: P300 (7257 NT)<br>* Gene 2: KDM5B (4647 NT)<br>* Frame interval: 5s<br>* Frames: 64<br>D: $0.21 pixels^2/s$<br>SNR 1- 36 |
| **$k_e mRNA_1$ vs $k_e mRNA_2$** | $k_e$: $2 - 12 aa \cdot s^{-1}$ | * Gene 1: P300 (7257 NT)<br>* Gene 2: KDM5B (4647 NT)<br>* Frame interval: 5s<br>* Frames: 64<br>* $k_i$: $0.06 s^{-1}$<br>D: $0.21 pixels^2/s$<br>SNR KDM5B: 2.6 - 14.2<br>SNR P300: 3.9 - 23 |
| **$k_i mRNA_1$ vs $k_i mRNA_2$** | $k_{initiation}$: $0.01 - 0.1 s^{-1}$ | * Gene 1: P300 (7257 NT)<br>* Gene 2: KDM5B (4647 NT)<br>* Frame interval: 5s<br>* Frames: 64<br>* $k_e$: $5.33 aa \cdot s^{-1}$<br>D: $0.21 pixels^2/s$<br>SNR KDM5B: 1.3 - 9.3<br>SNR P300: 1.7 - 13.3 |

**Figure 5:**
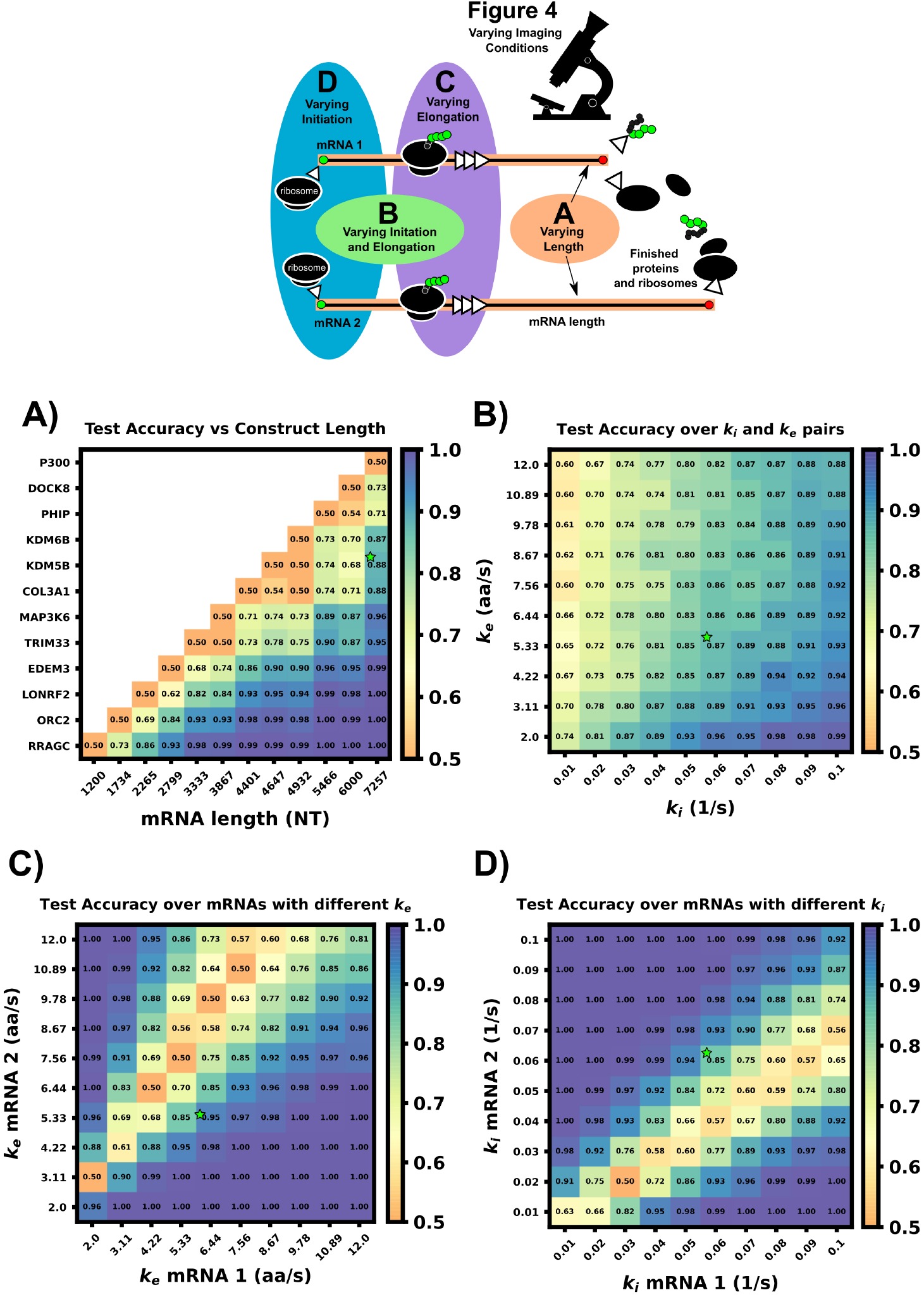
Comparison of ML test accuracy under variations in biophysical parameters. (Top) legend of which experimental parameters are changed for each panel. (A) Effect of construct length on classification test accuracy when trained on 4000 NCT spots and tested on 1000 withheld spots. Imaging conditions, initiation, and elongation are held constant while mRNA lengths are swept from 1200 NT to 7257 NT using different mRNAs. All classifiers are trained on 4000 NCT spots and tested on 1000 NCT spots to get the test accuracy. (B) Classification accuracy for P300 and KDM5B versus shared initiation and elongation rates. (C) Classification of P300 and KDM5B with shared initiation rate (0.06 1/s) but with different varying elongation rates. D) Classification of P300 and KDM5B with shared elongation rates (5.33 aa/s) and varying initiation rates. The green star in each panel denotes the default P300/KDM5B experiment with 5s frame interval, 64 frames, initiation rate of 0.06 1/s, and elongation rate of 5.33 aa/s.

### mRNAs with sufficiently different lengths can be differentiated using their fluctuation intensity signals

To explore the effect of mRNA length differences on classification, we applied our architecture to NCT experiments where the only difference between mRNAs is their length (Figures 5A and 6A). In addition to the previous P300 and KDM5B constructs, ten new genes with approximately evenly spaced nucleotide length coding regions were selected from the human consensus coding sequence database, Table 5 row 1. A standard 1011 nucleotide 10x-FLAG tag was added to the N-terminus end of each before simulating the NCT experiments. We assume a common set of global cell translation parameters, that is, every mRNA has a common ribosomal initiation and elongation rate since all mRNA would be in the same cell and they have been designed to have identical UTRs. Specifically, for this parameter sweep, we assume that *k*_e_ = 5.33 aa/s and *k*_i_ = 0.06 ribosomes/s and that video is recorded for a moderate length of 64 frames at a rate of one frame every five seconds. For each NCT simulation, the longer of the two mRNA species will retain ribosomes for longer periods of time and will therefore exhibit slower decorrelation times and higher average intensities.

For convenience, the difference between mRNA lengths can be quantified by the fold change:

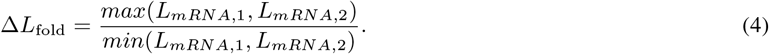

As one should expect, Figures5A and 6A show that as Δ*L*_fold_ becomes larger, the classification becomes easier for our machine learning architecture. For our simulated conditions and video length, we find that a 1.4-fold difference is sufficient to achieve greater than 80% classification accuracy of any two mRNA combinations of the 12 we selected. However, Figure 6A shows that one can lower the required length fold change by increasing the number of frames given. For example, if one extends imaging to 128 frames at 5s resolution (double the frames of the original condition), then one could achieve an 80% classification for a smaller Δ*L*_fold_=1.1. However, there is a diminishing benefit to adding extra video length; extending imaging to 1500 frames at 2s resolution (a barely achievable amount of data to collect with current NCT capabilities) provides only a marginal further improvement (compare teal and brown lines). This diminishing return emphasizes the need for careful consideration when designing NCT experiments with multiple mRNAs with identical tags, either to avoid designs that would require an unobtainable amount of sampling or to reduce sampling for designs that can be differentiated with fewer imaging resources.

**Figure 6:**
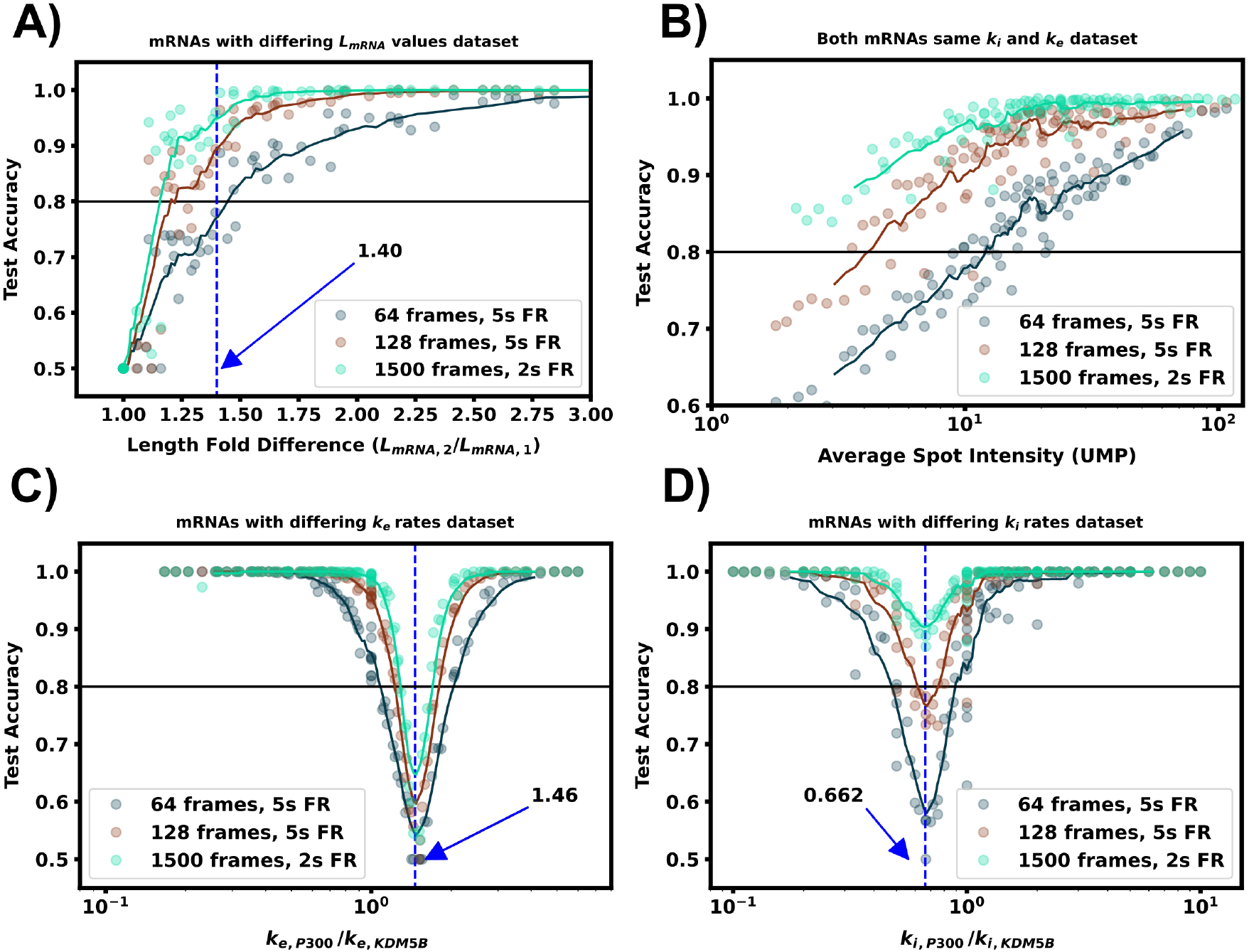
Increasing video length to resolve difficult to classify mRNA combinations. (A) Classification accuracy versus mRNA length fold difference, assuming identical tag designs and parameters and videos with 64, 128, or 1500 frames. (B) Classification accuracy for P300 and KDM5B with identical tags and parameters vs average P300 intensity (proxy for signal-to-noise ratio). As SNR, video length, and resolution increase, there is a corresponding increase in classification accuracy. (C) Classification accuracy versus ratio of P300 and KDM5B elongation rates. As parameters approach the dotted line at *k*_e,P300_/*k*_e,KDM5B_ = 1.46, the frequency and intensity information is identical between the two mRNAs, and increasing video length provides only marginal improvements. (D) Classification accuracy versus ratio of P300 and KDM5B initiation rates. As parameters approach the dotted line at *k*_i,P300_/*k*_i,KDM5B_ = 0.648, the two mRNA attain similar intensity means, but classification can be achieved through frequency content and is improved substantially by collecting longer videos.

### mRNA with different lengths can be distinguished for a range of different combinations of their biophysical parameters

In the next comparison, we sought to understand how classification accuracy depends upon the biophysical parameters that govern translation dynamics. Specifically, Figures 5B and 6B explore how the accuracy to classify P300 and KDM5B constructs would depend on their shared rates of translation initiation and elongation (*k_i_* and *k_e_*, respectively). Because the two constructs have fixed lengths in this comparison, every combination of (*k_i_, k_e_*) yields the same ratio of intensity mean and decorrelation time. Therefore, classification should be possible across the entire parameter range, provided that one obtains a sufficient level of video sampling. Figure 6B and Supplemental Figure S3B show that indeed, once there is enough video resolution, all (*k_e_, k_i_*) pairs can be classified with higher than 90% accuracy. However, when constrained to a fixed number of frames and frame interval (e.g., 64 frames with 5s frame interval as considered in Figure5B), some combinations of parameters yield signals that are brighter and are therefore easier to classify using intensity statistics. Specifically, when the initiation rate is high and the elongation rate is low, more ribosomes enter per second and remain longer on the mRNA. Conversely, “sparse” loading conditions prove harder to classify due to low ribosomal occupancy and rare ribosomal entry and thus, a lower signal to noise ratio.

In natural constructs, different 3’ and 5’ UTR sequences will affect the availability of initiation factors, and one should expect that translation initiation rates will vary from one mRNA species to the next (Gray and Wickens, 2003; Leppek et al., 2017; Fraser, 2015; Mayr, 2016). To explore how differences in initiation rate would affect classification accuracy, Figures 5C and 6C compare the classification accuracy as a function of both mRNAs’ unique initiation rates. In this case, it is possible for both mRNA to have very different or nearly identical intensity means depending on how close the ratio of the initiation rates compares to the inverse ratio of their lengths. For our particular case of KDM5B and P300 mRNA, if we neglect ribosomal collisions, the ratio of mean intensities can be estimated as (Aguilera et al., 2019):

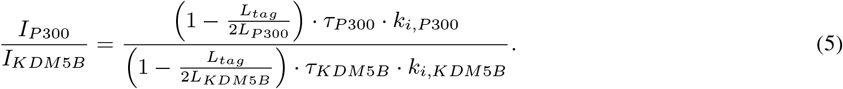

By setting this intensity ratio to unity, we can rearrange to find the corresponding ratio of initiation rates as

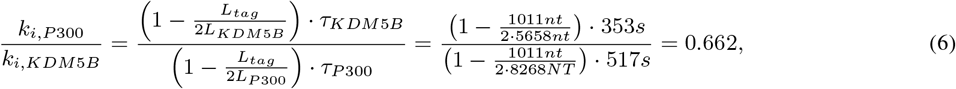

where we calculated the expected elongation times for the two mRNA (*τ*_KDM 5*B*_ and *τ*_*P* 300_) under an assumption of sparse loading for the ribosomes (i.e., no collisions). Figures 5C and 6C show that when the two mRNAs’ initiation rates approach this critical ratio, the accuracy decreases substantially. However, although these mRNAs may have similar intensities, their different lengths still result in distinct dwell times, and Figures 6C and S3 show that frequency information can still provide for accurate classification, especially as one increases the amount of video.

Figures 5D and 6D explore the opposite circumstance, where the two mRNA have the same initiation rate, but with two different elongation rates. In this case, it would be possible for both the intensity means and the dwell times to be identical for the two constructs if their elongation rates satisfy the ratio:

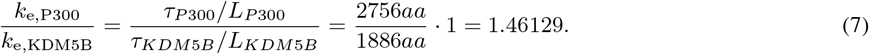

Figures 5D and 6D show that NCT signals along this parameter manifold are virtually indistinguishable as the only variation between their statistics is the time ribosomes spend in the tag region. Specifically, ribosomes on P300 reach full intensity fluorescence 1.46x faster than those on KDM5B. Increasing video resolution could potentially resolve parameter sets close to this manifold, but Figure 6D shows that accurate discrimination would require an unrealistic amount of NCT video. In conditions like these, it may be best to tag the mRNAs with different tag colors or use different tagging strategies as we discuss in the next section.

### mRNAs with similar fluctuations and intensities can be made more classifiable via intelligent design of tag placements

Considering the comparisons in Figure 5A-D, we observed that there are several conditions under which machine learning may be unable to classify NCT spots. Specifically, classification may fail when: (1) videos are too short to sample intensity distributions or have a too poorly chosen frame interval to quantify intensity frequencies (Figure 4); (2) when signal intensities are too low to capture the relevant statistics; or (3) when the mRNA lengths, initiation rates, or elongation rates combine such that both mRNAs yield identical statistics for both intensity and frequencies. As discussed above, the solution to the first two “failure modes” is to collect more information (i.e., longer videos with a more appropriate temporal resolution). In contrast, for conditions where intensity and frequency statistics are nearly identical, such as in Figures 5C and 6C, no amount of extra information or imaging will be able to tell these NCT spots apart. In such a circumstance, the similarity between the two mRNA could be ameliorated by changing the mRNA constructs themselves. For example, one could alter the fluorescent signal statistics either by lengthening one of the mRNAs in question or by employing an alternate tagging design. Changing the length of the mRNA with linker or junk regions may introduce unwanted effects in the mRNA / protein targets under study, so changing the tag region is preferable. To demonstrate this possibility, Figure 7A and B considers the case from above where the elongation rates of P300 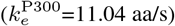 and KDM5B 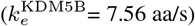 differ by the critical factor of 1.46, such that the 10xFLAG-P300 and 10xFLAG-KDM5B constructs yield identical intensity fluctuations that cannot be discriminated from one another. We then propose several modifications to the tagging scheme for KMD5B to explore how different designs might affect classification accuracy as follows:

- 10x Flag Tag on the N-terminus of KDM5B (original ineffective design)
- Splitting the tag region to relocate 3 epitopes to the C-terminus
- Relocating the tag region to the C-terminus of KDM5Bs CDS
- Adding 5 epitopes to the end of the 10x Flag Tag (adding in 5 ‘DYKDDDDK’ sequences separated by two glycines each)
- Removing 5 epitopes from the 10x Flag Tag (mutating the last 5 epitopes from ‘DYKDDDDK’ to ‘DYKDGGDK’)
- Relocating the 10x Flag Tag to the C-terminus of KDM5Bs CDS

**Figure 7:**
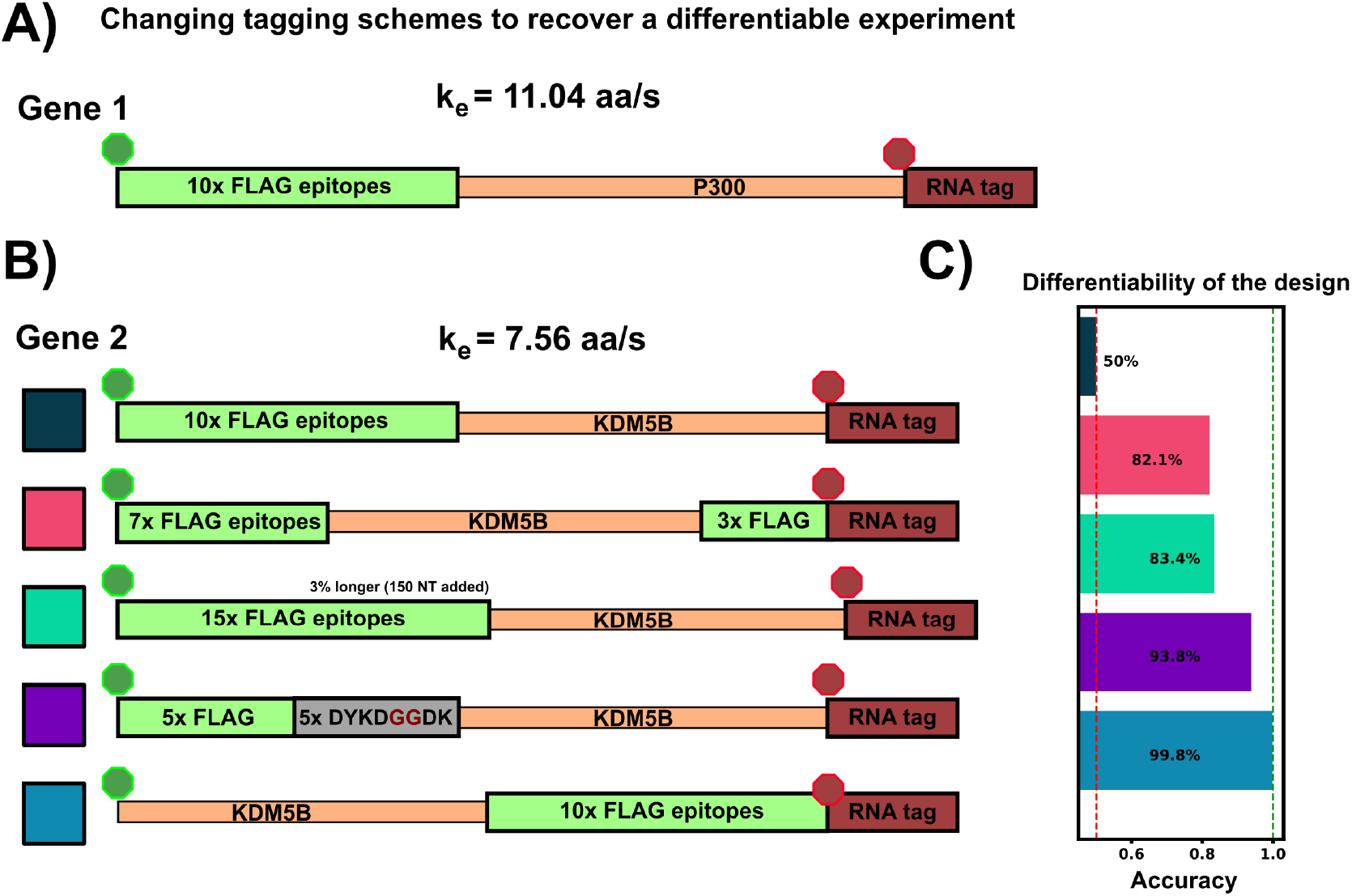
Changing tag designs to improve classification accuracy. (A) Tag design for P300 construct is kept fixed. (B) Five different tag designs for KDM5B created by splitting the tag, increasing or decreasing the amount of epitopes, or relocating the tag region to the 3’ end. (C) Accuracy for classification corresponding to each of the design combinations, and all assuming an elongation rate ratio of 1.46, under which the original design was non-classifiable (Figs 5D and 6D). All alternative designs would dramatically increase classification accuracy.

Figure 7(C) shows that any of these permutations to the original 10x Flag tag on the 5’ end of KDM5B allows the two mRNAs to be classified with greater than 85% test accuracy. Each tagging strategy changes the intensity dynamics of KDM5B spots, shifting them away from similar means and variances of the P300 spots, allowing classification without resorting to using a different color tag. One should note that each of these strategies comes with its own potential drawbacks: Moving the 10x flag to the end doesn’t allow any information about the upstream translation dynamics to be captured in the NCT experiment, and removing / moving 5 epitopes tag to the 3’ end creates a dimmer spot, potentially obscuring translation dynamics under study. Adding epitopes also has its own potential drawback of needing longer plasmids for transfection.

### Classifiers can retain accuracy despite uncertainties or assumption errors in biophysical parameters

In the previous sections, we explored how well classification would work in the ideal situation in which the classifier is trained on data that match (in probability) to the circumstances of the testing data (although every intensity trajectory is different due to stochastic fluctuations in translation, motion, and cellular background noise). In other words, the stochastic model used to generate the training data was the same as the model used to generate the testing data. For a more realistic test of how well one might expect a classifier trained using simulated data to work when applied to experimental data, one must acknowledge that true biophysical parameters are unknown, and they may vary from cell to cell or from one individual mRNA to the next. For example, in the analysis of KDM5B and P300, it is reasonable to assume that the mRNA designs and lengths are known, but one might only have a rough estimate for the initiation and elongation rates based on analyses for other mRNA, cells, or conditions. Ideally, the classifier should still work despite finite errors in these parameter estimates. To explore how well a simulation-trained classifier might work when parameters are incorrect, Figure 8 shows the accuracy versus *unknown* rates *k_e_* and *k_i_* when the model is trained at three specific assumptions for those rates (denoted by red squares in Figure 8). When the model is trained with a fast elongation rate and a slow initiation rate (Figure 8A), classification accuracy is always poor (as discussed above, see Figure 5B). However, when the model is trained in a condition that is more conducive for accurate classification (Figure 8B), the accuracy is strong not only for the exact parameters under which the model was trained, but for a large range of surrounding parameter sets. The practical implication of this result is that one could in principle use an approximate model (e.g., the simulations presented in this study) to train a classifier and then reliably trust that classifier despite unavoidable but finite errors in the underlying model or parameter assumptions. To maximize the utility of such a classifier, we performed a search over all possible sets of parameters on which to train the model and asked which training set leads to a classifier that would work best when averaged over the unknown “true” parameters. Figure 8C) depicts the accuracy of this model (which is trained at *k_i_* = 0.07 s^−1^ and *k_e_* = 6.44 aa/s) as a function of all parameters, and it results in an expected average classification accuracy of 70% but with classification greater than 80% for large regions of parameter spaces.

**Figure 8:**
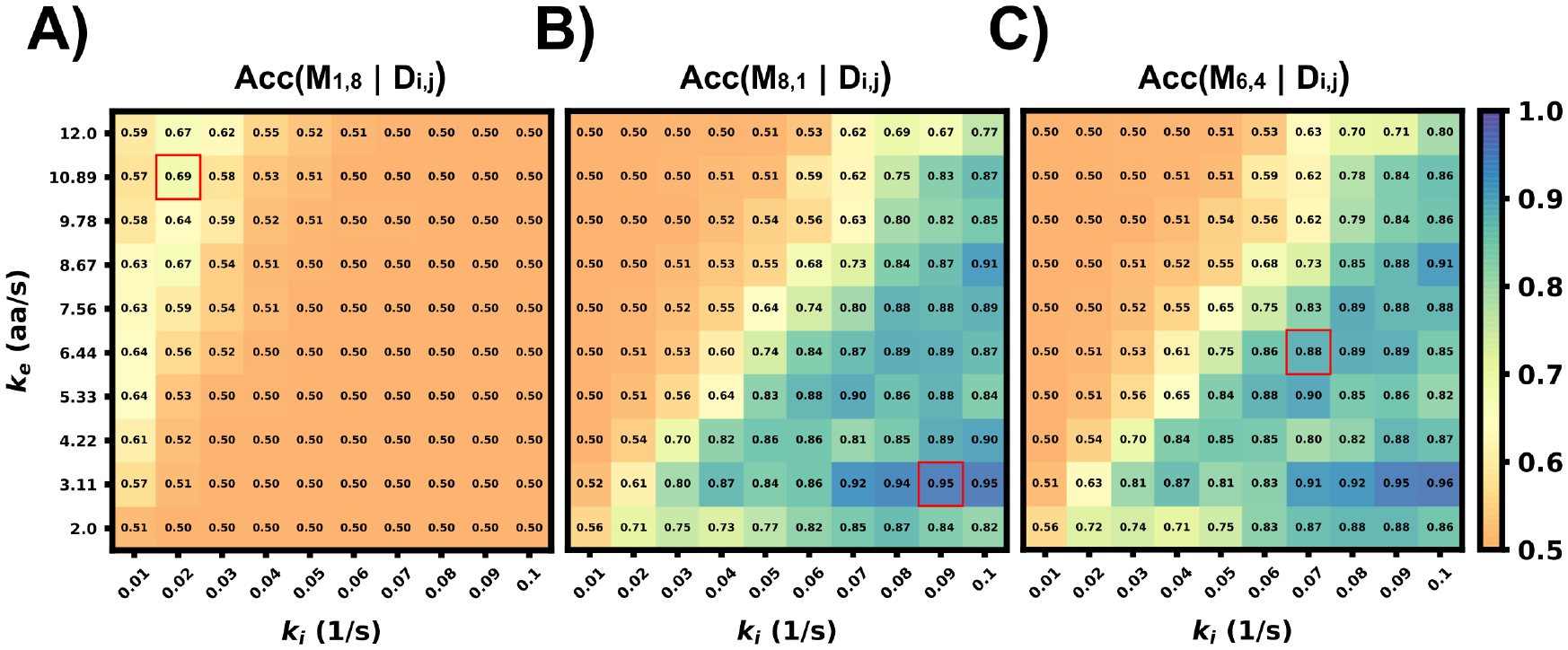
Accuracy of classifier when trained with incorrect parameter assumptions. Accuracy versus the actual rates *k*_e_ and *k*_i_ when the model is exclusively trained on three specific, but possibly incorrect, sets of these parameters: (A) (*k*_i_ = 0.02s^−1^, *k*_e_ = 10.89*aa*/s), Average accuracy = 52.4%, (B) (*k*_i_ = 0.09s^−1^, *k*_e_ = 3.11*aa*/s), Average accuracy = 70.1%, (C) (*k*_i_ = 0.07s^−1^, *k*_e_ = 6.44*aa*/s), Average accuracy = 70.2%,

### Using combinations of intensity fluctuation information and different color mRNA tags, one could design experiments to distinguish several mRNAs within the same cell

Finally, to highlight the how our proposed pipeline might be used to increase the potential of NCT experiments, we demonstrate it on the simulation of seven tagged species within a single cell. Using the Figure 5A heatmap, we selected four mRNAs species that were differentiable from each other with a higher than 90% accuracy for the green channel, and three mRNA species for the blue channel with the same accuracy threshold, Table 6 and Table 6. A single simulated “multiplex” cell video was generated with the conditions described in both tables. Ten spots for each of the seven mRNAs were added to the appropriate color channel for each cell.

**Table 6:** Multiplexing simulation parameters

| Comparison Analysis | Variable Range | Constants |
| --- | --- | --- |
| **Green Channel (4 spots)** | mRNA length (without tag):<br>* RRAGC (1200 NT)<br>* LONRF2 (2265 NT)<br>* MAP3K6 (3867 NT)<br>* DOCK8 (6000 NT) | * Frame interval: 5s<br>* Frames: 64<br>* $k_i$: $0.06 s^{-1}$<br>* $k_e$: $5.33 aa \cdot s^{-1}$<br>D: $0.925 \mu m^2/s$<br>SNR: 3.7-15.2 |
| **Blue Channel (3 spots)** | mRNA length (without tag):<br>* ORC (1734 NT)<br>* TRIM33 (3333 NT)<br>* PHIP (5466 NT) | * Frame interval: 5s<br>* Frames: 64<br>* $k_i$: $0.06 s^{-1}$<br>* $k_e$: $5.33 aa \cdot s^{-1}$<br>D: $0.925 \mu m^2/s$<br>SNR: 5.2-14.9 |

An additional class of non-translating spots was also added by taking 2500 trajectories from the opposite (no-spot) channel, but with the same Brownian motion. The ML architecture was adjusted to account for the multiple mRNA labels and the noise label for non-translating spots, and the final layer was set to a softmax layer and an output of the number of species in each color channel (four for blue and five for green, including one the non-translating spots in each channel). A model was trained with the process described in the ML section for each color channel using the matching data from the construct length dataset, (7500 NCT spots for blue, 10000 spots for green with 80:20 train-test split and three-fold cross validation). Test Video trajectories were normalized with the same min-max scaling as the training data.

After training, the models were applied to classify the simulated trajectories in 50 new multiplexing videos. Figure 9A shows representative images of the artificial ML labeling results applied to an example multiplexing video. Correctly identified spots are denoted by circles and incorrect classification results are shown with ’x’. A representative crop for each spot type is shown on the left. Figure 9B shows the confusion matrix for each mRNA class and the blank trajectories. The green channel classifier had an 81% accuracy (disregarding blank trajectories) and struggled mostly with the two middle length genes LONRF2 and MAP3K6. The blue channel classifier had a 91% accuracy disregarding noise only spots. As expected, the majority of misclassified spots were to their length neighbors that have the most similar statistics. The shortest, dimmest mRNA NCT spots, ORC2 and RRAGC, were still classifiable from the non-translating spots with almost 100% accuracy. 2.2% of RRAGC spots were misclassified as noise. Overall on the test video, the classifier correctly identified 64 out of 70 spots, demonstrating the potential to label multiple species in the same cell with a correct tagging scheme and ML labeling.

**Figure 9:**
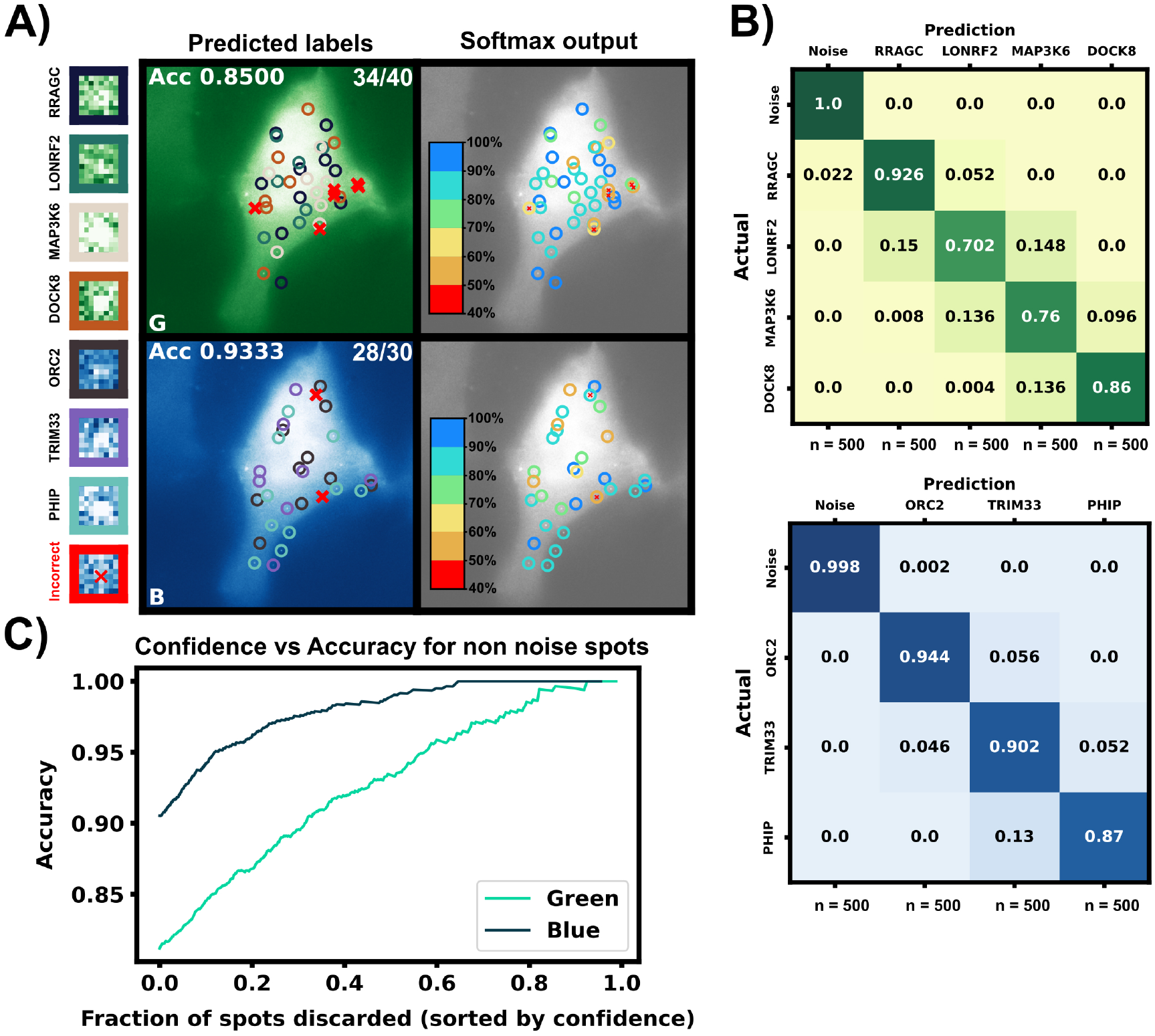
Simulated multiplexing of seven different mRNA species in a single cell. Ten mRNAs each of RRAGC, LONRF2, MAP3K6, and DOCK8 with identical were simulated in the green channel, and ten mRNAs each of ORC2, TRIM33, and PHIP were simulated in the blue channel with our pipeline, and all with identical tag designs and parameters. Our architecture was modified for multiclass labeling, and a model was trained for the green and blue channel for artificial labeling of the example video. (A) Example frame from video classification with seven different mRNA transcript types. Incorrectly labeled spots are marked with an X (6/70 spots). Crops of example spots are show to the left. (B) Confusion matrices for the green and blue channels when tested on 50 cells containing 10 spots of each mRNA. (C) Accuracy of the classifier versus the fraction of low-confidence spots that is discarded. If one only considers the 50% most confident spots, then accuracy rises to 93.4% and 98.9% for the blue and green channels, respectively.

In addition to the discrete labels generated by the classifier, the softmax output (shown in Figure 9C, left) provides a quantification of the classification confidence for each spot. Figure 9C shows the effect of sorting all classified non-noise spots by their softmax output and discarding those spots with the lowest confidence for classification. By discarding 50% of the spots with the lowest confidences yields a dramatic improvement in accuracy for both the green channel (from 81% to 92%) and blue channels (from 90% to 98%). In other words, although perfect classification may not be achievable, one can focus on confidently identified mRNA to analyse their behaviors, e.g., to determine how different mRNA types respond to subsequent cellular signals, drugs, or stress perturbations. It is important to note, our architecture is small but no softmax calibration was used; a better metric of confidence could potentially be obtained by using a softmax calibration in future works.

## 4 Conclusion and Future Work

Temporally- or spatially-resolved activation or repression of translation provides for a potent mechanism by which cells could rapidly alter their protein content in response to cellular signals (Sonenberg and Hinnebusch, 2009; Fabian et al., 2010; Hershey et al., 2012; Zhao et al., 2019; Peer et al., 2019). Recent advances in Nascent Chain Tracking (NCT) experiments has made it possible to observe this regulation at the level of single mRNA molecules in living cells (Lyon et al., 2019; Moon et al., 2019; Koch et al., 2020; Cialek et al., 2022; Kobayashi and Singer, 2022). However, limitations on the number of distinct fluorophores prevents current NCT experiments from exploring more than one or two different mRNA species at a time.

In this work, we use computational simulations to propose a solution to circumvent this limitation. Specifically, we provide a pipeline (Fig. 1) to combine mechanistic models (including detailed simulation of nascent protein elongation and corrupted by fluorescence background and camera noise) with machine learning to classify mRNA species based on their fluctuating fluorescence intensity signals in NCT experiments. We show that multiple mRNAs labeled with identical fluorescent tags could be distinguished provided that the mRNAs have some variation in their intensity distributions or fluctuation frequency content, e.g., due to different lengths (Figs. 5a, 6a) or different translation parameters (Figs. 5B-D). We also demonstrate how our computational pipeline could help to guide the design of experiments to make it easier to access these features and distinguish between different types of translating mRNA. Specifically, by changing multiple biophysical parameters or experimental design variables – e.g., changing tag design (Fig. 7), mRNA lengths (Fig. 5a), ribosomal elongation and initiation rates (Figs. 5 B-D), or the length and temporal resolution of NCT movies (Fig. 4) – we explored which realistic designs of NCT experiments would provide insight for classification, and which would not allow for NCT multiplexing.

Our realistic simulations show that that ML labeling accuracy higher than 80% can be achieved under reasonable NCT experimental settings (Figs. 2-9). Longer videos with appropriately-chosen frame intervals would lead to a better classification of NCT signals, albeit with diminishing returns (Figs. 4,6). A more strategic approach shows that selectively tagging pairs of mRNA species to achieve the greatest difference in expected intensity or frequency content achieves higher classification with a minimal number of frames (Fig. 9). We also demonstrated that different tagging strategies (Fig. 7) can help to separate hard to classify mRNA species, with tag design options ranging from simply adding more tag epitopes to increase one mRNAs intensity, to relocating or dividing the tag region between the 5 and 3 end of the CDS to alter the frequency content. Using these strategies, additional fluorophore colors can be held in reserve for mRNAs that are too similar in their lengths or dynamics, and we demonstrate that our multiplexing pipeline could distinguish seven different mRNAs at greater than 90% accuracy when using only two different fluorescent tag colors and only one per gene (Fig. 9). Additional mRNA could be considered by combining multiple colors on the same mRNA (e.g., mRNA with red and green could easily be distinguished from those with red or green alone).

Moreover, to evaluate the potential to transfer our model-based findings to real experiments, we verified that models learned from one set of mechanistic parameters could correctly classify mRNA when tested on data that are generated using different parameters guesses (Fig. 8). This fact that classification accuracy remains high despite inexact knowledge of the model parameters offers hope that classifiers learned using approximate models could work on data from real experiments, without a need for collecting excessive training data. Based on these promising results of our detailed simulations, we envision that the next generation of NCT experiments will be able to track and differentiate multiple identically-tagged translating mRNA within the same cells (especially if these use the identified experiment designs for tag positions, gene lengths, and video frame interval).

Although, for the sake of brevity, this paper does not explore all mechanisms that can be analyzed by the rSNAPsim and rSNAPed computational pipeline (e.g., effects of tracking, photo-bleaching, variable probe-biding rates, ribosome pausing, etc.), future work could expand on these capabilities along with further exploration of different machine learning approaches (e.g., different ML architectures or different types of classifiers) or inclusion of additional features (e.g., including mRNA diffusion rates, cell position information, fluorophore bleaching rates, etc.). Similarly, although the current manuscript has focused exclusively on supervised learning techniques that require known ground truth (e.g., labels in simulated data), one could potentially improve the application to real data through the addition of unsupervised machine learning or transfer learning approaches. For example, the simulation-based pipeline proposed here could be used to design tags and experimental conditions, while unsupervised approaches may be applied to differentiate spots in subsequent experiments. Finally, beyond the goal of multiplexing, we believe that our general approach to combine detailed mechanistic models, realistic simulations of microscopy and image analyses effects, and machine learning classifiers could help to design improved experiments that are more suited to other biological questions (e.g., to differentiate between competing hypotheses for translation mechanisms rather than to differentiate different mRNA species as explored here).

## Author Contributions

Conceptualization: ZRF, TM, ER, TJS, BM. Theory and computational modeling: WSR, LUA, ZRF, MPM, BM. Computational Simulation: WSR, LUA. Machine learning: WSR, SG. Writing: WSR, BM. Editing: WSR, LUA, SG, TM, TJS, BM. Resources, supervision, and funding acquisition: TJS, BM.

## Acknowledgements

This manuscript has been authored by UT-Battelle, LLC under Contract No. DE-AC05-00OR22725 with the U.S. Department of Energy. The United States Government retains and the publisher, by accepting the article for publication, acknowledges that the United States Government retains a non-exclusive, paid-up, irrevocable, world-wide license to publish or reproduce the published form of this manuscript, or allow others to do so, for United States Government purposes. The Department of Energy will provide public access to these results of federally sponsored research in accordance with the DOE Public Access Plan (http://energy.gov/downloads/doe-public-access-plan).

## Data Availability Statement

The datasets generated and analyzed for this study can be found in the MunskyGroup/Multiplexing project Github Repository https://github.com/MunskyGroup/Multiplexing_project. Large files (saved classifiers, example videos, dataset CSVs) are stored at: https://www.dropbox.com/scl/fo/cj4fd2o0nr5dfstngyh83/h?dl=0&rlkey=363ka3o1ur8jvywhh5mvolxqg.

---

**Figure 1:**
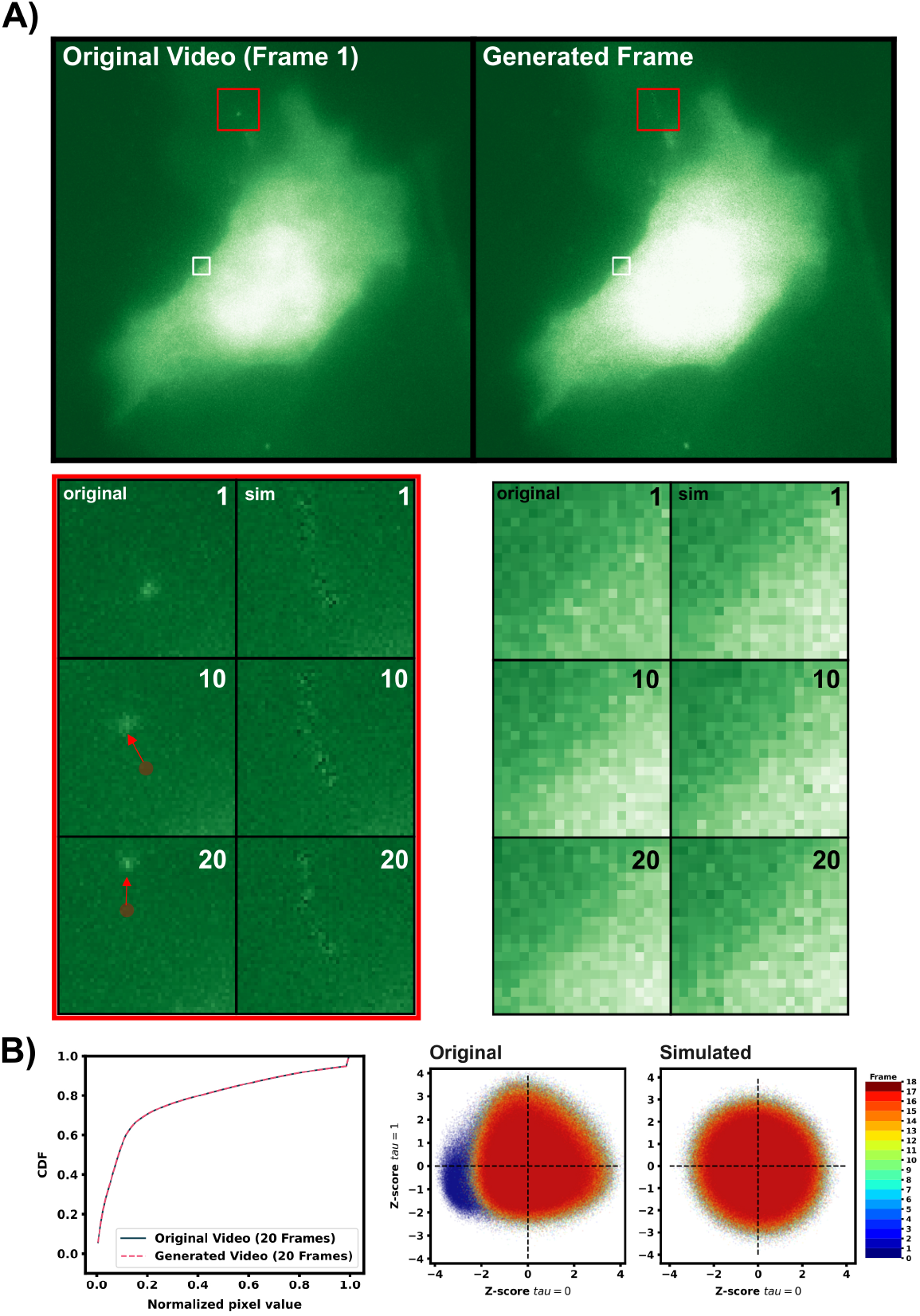
Gaussian generation of simulated cell backgrounds. Each pixel within the original blank cell video (20 frames) is fit to its own Gaussian distribution. New frames are generated pixel by pixel by pulling new values from that array of Gaussians. Standard deviations above the 95th quantile of the total videos per pixel standard deviation are set to that 95th quantile value that way problem pixels that may be dark for 19 out of 20 frames do not result in a massively wide Gaussian to pull from. For the purposes of visualization, videos were min max normalized by the 95th quantile intensity and the 1st quantile value of the original video. The red square highlights a region where a large moving feature in the original video is lost by the Gaussian generation approach. Large bright features that move over the course of 20 frames are represented as a larger standard deviation and a slightly higher Gaussian mean, resulting in a smeared “noisy region in the generated videos that overlays the course of the original bright feature. The white box shows an example highlighted area with no moving feature. Below shows the CDF of each videos histogram of normalized pixel values. We note that for all 7 background videos the first frame is noticeably dimmer than all the rest (blue region in panel B middle).

**Figure 2:**
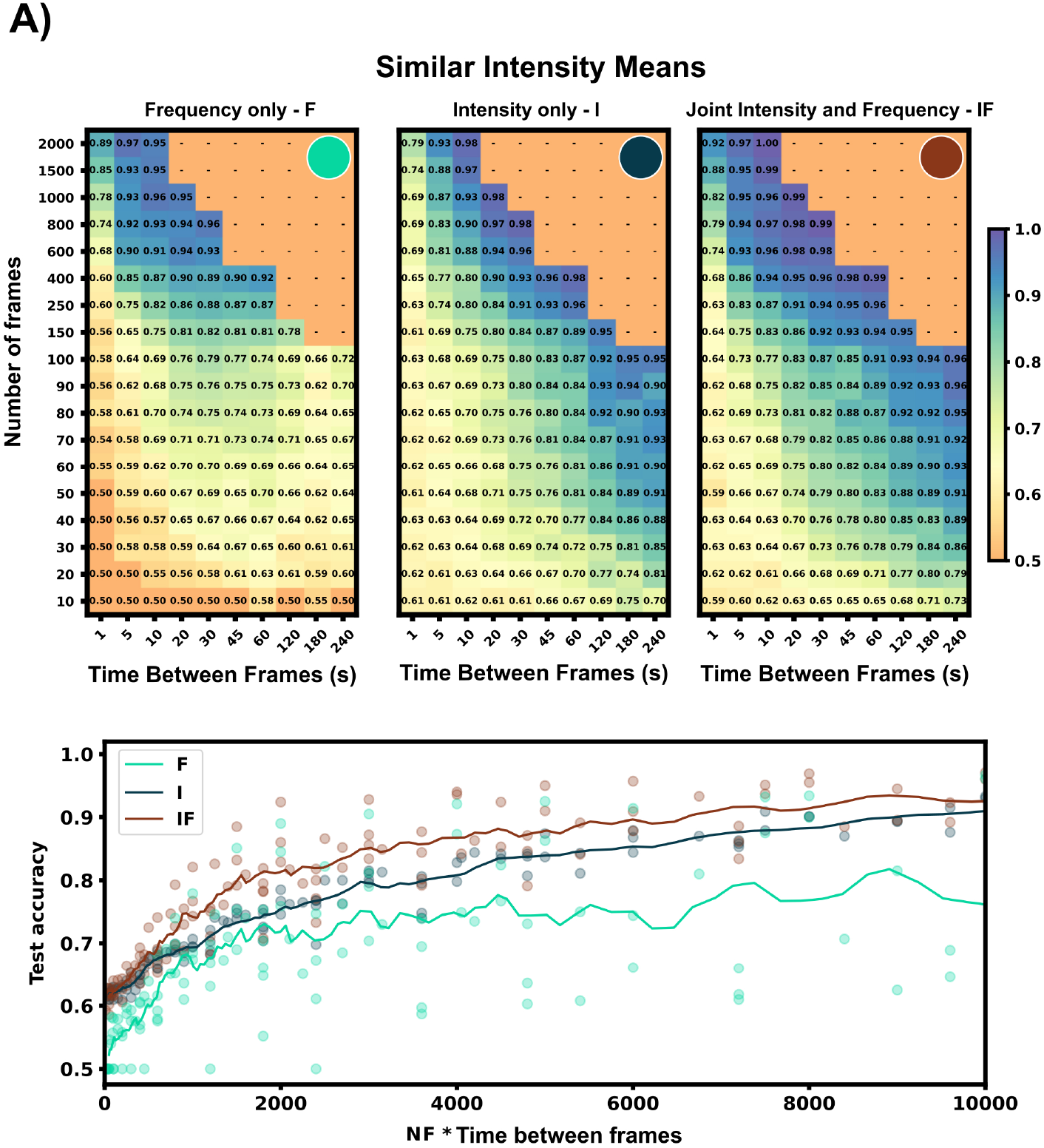
(Top) Accuracy of separate architecture halves (I, F) and joint architecture (IF) across a large range of frame rates and number of frames. Heat maps show each architecture half or the joint architecture applied on a NCT data set with different frequency content (decorrelation times, *τ* = 354s vs *τ* =517s) and similar intensity means (4.7 UMP vs 6 UMP). (Bottom) Using both architecture halves increases average test accuracy across all conditions vs the individual feature architectures. All test accuracies from the heat maps above are plotted vs their video length (NF * Time between frames) for the full architecture (IF), and each half architecture (I, F). A trend-line was generated with a moving average of 100 seconds.

**Figure 3:**
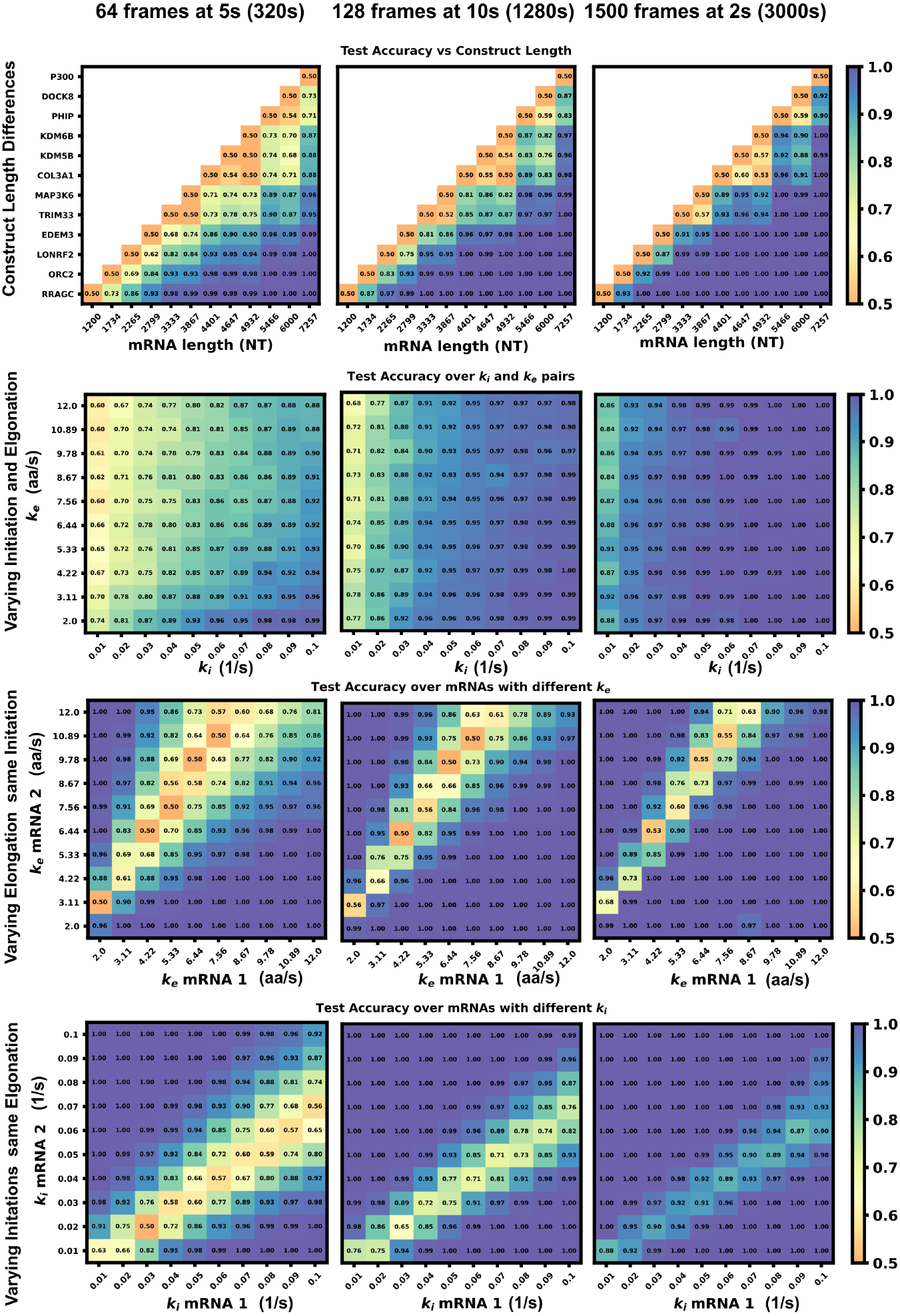
Machine learning accuracy for increased video size and resolution. Panels on left column are duplicated from Figure 5A-D in the main text (64 frames at 5s frame interval) and compared to the accuracy with increased video size of (middle) 128 frames at 10 s frame interval and (right) (1500 frames at 2 second frame interval)

## Footnotes

1 Problem pixels that have extremely large standard deviations (e.g., for a video where a given pixel’s intensity becomes extremely bright for one randomly-timed frame) are corrected to the 95th percentile of all pixels’ standard deviations.

